# An autologous human iPSC-derived 3D organoid infection model for preclinical testing of antiviral T cells

**DOI:** 10.1101/2025.09.16.676521

**Authors:** Ugarit Daher, Valeria Fernandez-Vallone, Morris Baumgardt, Benedikt Obermayer, Niklas Wiese, Achim Klaus Kirsch, Tanja Fisch, Anna Löwa, Michael Schmueck-Henneresse, Andreas C. Hocke, Leila Amini, Harald Stachelscheid

## Abstract

Immunocompromised patients, such as those undergoing hematopoietic stem cell or solid organ transplantation, are highly susceptible to viral complications. Given the limitations and side effects of available antiviral therapies, adoptive transfer of antiviral T cells offers a promising alternative by restoring immune defense. However, existing models for evaluating antiviral T cell therapies lack physiological relevance, limiting accurate predictions of efficacy and safety. There is a critical need for *in vitro* human infection platforms that support personalized assessment of therapeutic responses. To address this, we developed antiviral T cell products (TCPs) targeting Influenza A virus (IAV)-infected cells, alongside an autologous human induced pluripotent stem cell (iPSC)-derived 3D lung organoid infection platform. This model recapitulates key immunological responses and is compatible with a new 3D high-throughput, high-content imaging pipeline. Our study provides the first proof-of-concept for assessing T cell-mediated cytotoxicity in a 3D *in vitro* lung infection model, advancing personalized antiviral immunotherapy development.

## Introduction

Immunocompromised patients, including those that underwent hematopoietic stem cell (HSCT) or solid organ transplantation (SOT) often suffer from viral complications, accounting for about 40% of transplant-related mortality ^1^. Infections with cytomegalovirus (CMV), Epstein-Barr virus (EBV) or respiratory viruses such as adenovirus (ADV), severe acute respiratory syndrome coronavirus type 2 (SARS-CoV-2) or influenza A virus (IAV) are among the most prevalent causing significant morbidities and deaths ^1–5^. Adverse effects, resistances, poor efficacy or the complete lack of available antiviral treatments underline the need for novel therapeutic modalities ^6^. The adoptive transfer of antiviral T cell products (TCPs) represents a promising avenue in both the prevention and fight against viral infections by leveraging the patient’s own immune system to specifically target and eliminate virus-infected cells. Although antiviral TCPs have already shown promising results in clinical settings in-depth knowledge about their safety and efficacy profiles and concrete mode of action are lacking ^7–14^. Thus, preclinical functional testing is crucial for improving the selection of adequate cellular therapies and accelerating their translation into the clinics. The most widely used infection models are either animal models or 2D-based cell cultures ^15^. Although humanized animals represent the most complex available model, they are not suitable as preclinical infection platforms, because they poorly reflect human pathogenesis. In most cases, animal models lack the molecular repertoire to address the natural host of the pathogen and fail to reflect species-dependent immune responses ^16–18^. Furthermore, humanized animal models are not always readily available and require considerable time to be generated after the emergence of a novel pathogen, a limitation which became particularly evident during the COVID-19 pandemic ^16,17^. Human 2D cell culture models using immortalized cell lines or singularized organoids offer advantages in terms of reproducibility, ease of control and cost effectiveness ^19–22^. However, they often fail to accurately replicate the physiological setting of viral propagation and cell-cell interactions which are particularly relevant in infection studies ^20,22,23^. This highlights the need for more predictive models that bridge the limitations between traditional animal models and 2D-based human cell cultures. Organoids represent a promising avenue by providing a more complex, multicellular environment that better mimics aspects of human tissue function and architecture ^24^. In recent years, an increasing number of studies have paved the way for using human organoids as preclinical test platforms including screening of chemical compounds, biologics or even early-stage testing of cellular therapies such as chimeric antigen receptor (CAR) T cells in the oncology field ^25–31^. However, the use of 3D *in vitro* lung infection models for preclinical safety and efficacy assessment of virus-specific TCPs has not been systematically evaluated.

In this study, we present a good manufacturing practice (GMP)-compliant workflow for generating antiviral TCPs targeting IAV-subtypes H3N2 and H1N1. These products were phenotypically characterized and shown to specifically eliminate viral peptide-loaded antigen presenting cells (APCs). To model infection and evaluate T cell efficacy in a physiologically relevant system, we generated autologous human induced pluripotent stem cell (iPSC)-derived alveolar-like lung organoids (iPSC-aLOs). Transcriptomic analysis confirmed pulmonary lineage commitment in these organoids, which despite representing an early stage of pulmonary development, express key immunological markers such as HLA class I, essential for interactions with CD8^+^ T cells. Upon IAV challenge, we evaluated viral response of the iPSC-aLOs and confirmed induction of antiviral pathways. Furthermore, we established an automated high-throughput high-content imaging pipeline to study the interactions between IAV-infected iPSC-aLOs and autologous IAV-specific TCPs (IAV-TCPs) at single-cell resolution within a 3D context. Our results indicate that this approach enables prediction of TCP-mediated specific elimination of virus-infected cells within the organoids. We propose that our testing platform represents a significant advancement towards more predictive preclinical pipelines for safety and efficacy evaluation and selection of candidate therapies, thus facilitating the developmental process of personalized immunotherapies.

## Results

### GMP-compliant manufacturing of IAV-TCPs

To generate IAV-TCPs for the H3N2 and H1N1 subtypes, we adapted a GMP-compliant protocol to the G-Rex6M culture platform ^32–34^. The manufacturing process involved the isolation of virus-specific T cells from small volumes of peripheral blood mononuclear cells (PBMCs) based on their interferon-gamma (IFN-γ) production following stimulation with overlapping peptide pools (Figure 1A). Based on previous studies using inactivated viruses or peptides for IAV-specific T cell stimulation, we combined peptide pools of variable and conserved proteins, to target a broad antigen repertoire to yield a CD4^+^ and CD8^+^-balanced TCP ^35–37^. Hence, we combined pools covering the sequences from the internal nucleoprotein (NP) and matrix protein (MP) as well as the external protein hemagglutinin (HA). PBMCs were accordingly stimulated with either H3N2_HA/NP1/MP_ or H1N1_MP1/MP2/NP/HA_ to generate the respective TCPs. To assess the robustness and reproducibility of our protocol, we conducted multiple independent production rounds for both H3N2-and H1N1-specific TCPs (H3N2-TCPs and H1N1-TCPs). In total, we generated 17 batches of H3N2-TCPs from six different donors and 13 batches of H1N1-TCPs from four donors. To determine the purity of IFN-γ-sorted cells, we analyzed the frequency of antigen-reactive T cells pre-and post-enrichment (Figure 1B+C). Inter-experimental variability was observed for the same donors, with differences in post-enrichment purities between independent manufacturing rounds (%CD3^+^IFN-γ^+^ cells Donor 5: H1N1=23.18 ± 21.67 n=6, H3N2=31.83 ± 13.65 n=5; Donor 6: H1N1=20.29 ± 14.57 n=4, H3N2=33.31 ± 23.50 n=4) for both H3N2-and H1N1-TCPs (Figure 1C).

**Figure 1.**
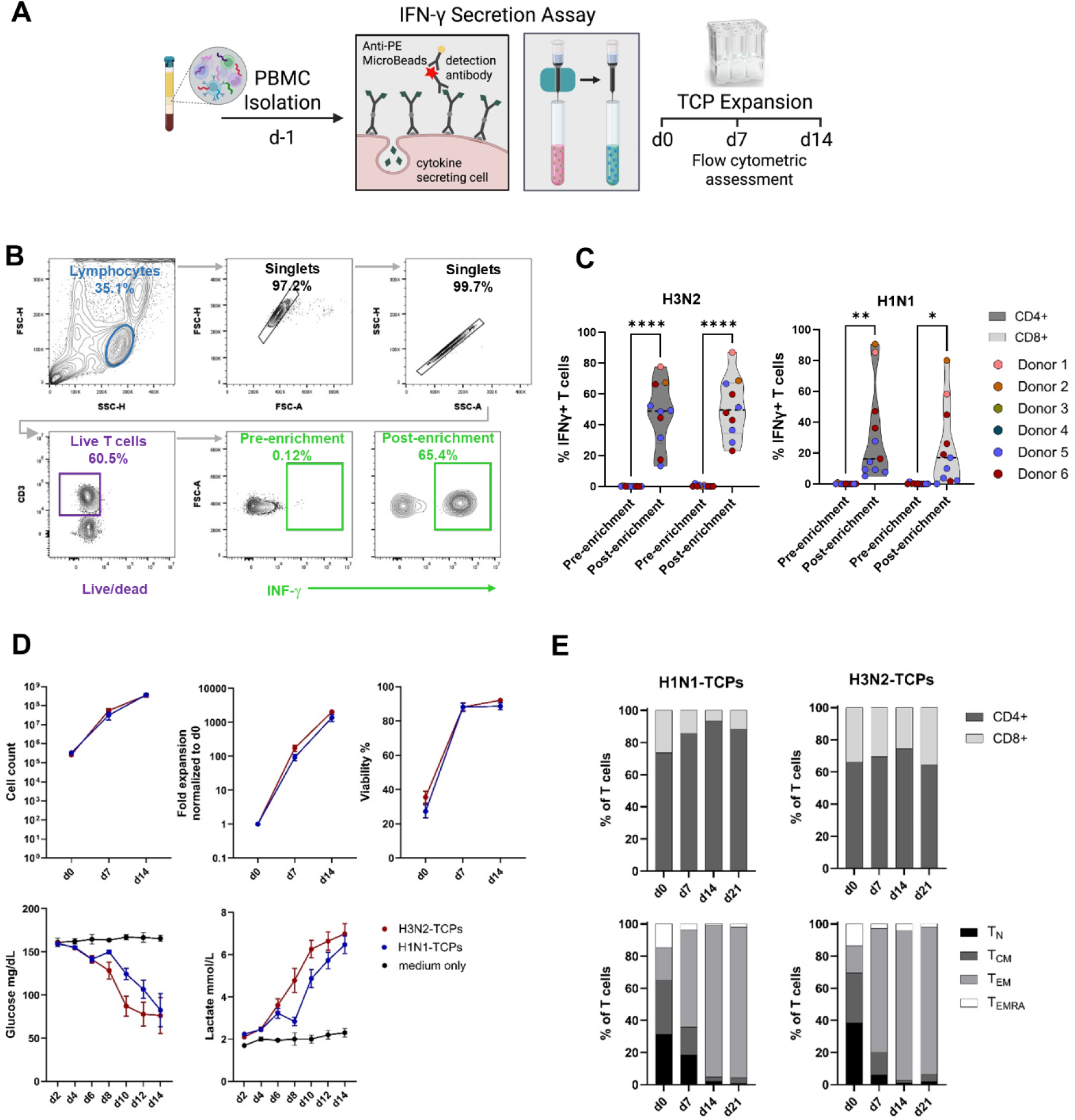
GMP-compliant manufacturing of IAV-specific TCPs. **A)** Scheme of GMP-compliant manufacturing process of IAV-TCPs. The image was created with BioRender.com. **B)** Representative flow cytometric plots showing gating strategy for assessment of pre-and post-enrichment of IFN-γ^+^ T cells. **C)** Violine plots showing frequencies of CD4^+^ and CD8^+^ IFN-γ^+^ T cells of pre-and post-H3N2 and H1N1 enrichment, respectively. Two-way ANOVA p<0.05 (*), p<0.01 (**), p<0.001 (***), p<0.0001 (****). **D)** Plots showing cell count, fold expansion, cell viability and glucose and lactate concentrations in culture medium over the course of expansion of H3N2-and H1N1-TCPs. Mean ± SEM (n=8-16, medium control n=2). **E)** Flow cytometric phenotype determination of H3N2 (n=17) and H1N1 (n=13) IAV-TCPs at indicated timepoints for CD4^+^ and CD8^+^ T cells. Upper panel: ratios of live CD4^+^/CD8^+^ T cells. Lower panel: composition of T cell pool with differentiation subsets CCR7^+^/CD45RA^+^ naive T cells (T_N_), CCR7^+^/CD45RA^-^ central memory T cells (T_CM_), CCR7^-^/CD45RA^-^ effector memory T cells (T_EM_), and CCR7^-^/CD45RA^+^ terminally differentiated effector memory T cells (T_EMRA_).

Post-enrichment cell viability ranged between 20-40%. Nevertheless, the G-Rex6M culture system supported significant expansion and recovery of IAV-TCPs, yielding clinically relevant cell numbers after 14 days of expansion for both specificities (Figure 1D) ^38,39^. Since visual inspection possibilities of the cells cultured in the G-Rex6M system are reduced, we implemented metabolic monitoring by tracking glucose consumption and lactate production as surrogates of cell proliferation and metabolic activity (Figure 1D). Media exchange was performed when glucose levels dropped below 50 mg/dL or when lactate concentrations exceeded 7 mmol/L. Phenotypic analysis of IAV-TCPs at key time points (day 0, 7, and 14) revealed a CD4^+^ dominated T cell population, which was consistently maintained throughout the entire expansion period for both H3N2-and H1N1-TCPs (Figure 1E). The CD4^+^/CD8^+^ ratio remained stable across batches and donors, with CD4^+^ T cells comprising between 60-90% of the final T cell population.

Key surface markers associated with T cell memory subsets were assessed to characterize the differentiation status of the expanded IAV-TCPs. The following subsets were defined in descending order of differentiation: CCR7^+^/CD45RA^+^ naive T cells (T_N_), CCR7^+^/CD45RA^-^ central memory T cells (T_CM_), CCR7^-^/CD45RA^-^ effector memory T cells (T_EM_), and CCR7^-^/CD45RA^+^ terminally differentiated effector memory T cells (T_EMRA_). IAV-TCPs post-enrichment were predominantly composed of T_N_ and T_CM_ subsets, indicating that the starting population consisted mainly of less differentiated T cells (Figure 1E). However, within the first week of *in vitro* expansion, a marked shift in subset composition was observed, with a substantial reduction in T_N_ and T_CM_ frequencies. By day 7, the T_EM_ phenotype remained the dominant subset throughout the expansion period. By day 14, T_EM_ accounted for approximately 70–90% of the final TCPs.

### IAV-TCPs display effector functions *in vitro* and eliminate target-specific cells

To evaluate the functional characteristics of IAV-TCPs throughout the expansion period, flow cytometric analysis was conducted following antigen-specific re-stimulation (Figure 2A and Supplementary Figure 1A). IAV-TCPs demonstrated a robust upregulation of the activation markers CD154 and CD137, alongside increased production of IFN-γ, IL-2 and TNF-a at both day 7 and day 14 (Figure 2 A,B). Notably, under the applied culture conditions, granzyme B (GrzB) was pre-formed (Figure 2A), making it the only marker for which background subtraction from unstimulated controls was not performed. IAV-TCPs retained their capacity for antigen recognition and activation throughout the expansion phase. H3N2-TCPs exhibited a slightly higher degree of cytokine production and activation marker expression, suggesting a more pronounced response compared to H1N1-TCPs (Figure 2B). A time-course analysis of IFN-γ revealed that both CD4⁺ and CD8⁺ T cells contributed equally to its production early in expansion. IFN-γ levels declined between day 0 and day 7, then stabilized through day 14, indicating a decrease followed by sustained cytokine output. In contrast, IL-2 levels remained relatively stable throughout, with no significant differences between subsets. TNF-α production showed subset-specific dynamics, with CD8⁺ T cells maintaining stable levels and CD4⁺ T cells exhibiting a significant decline between days 7 and 14. GrzB was shown to be predominantly produced by CD8⁺ T cells, consistent with their cytotoxic function. Finally, expression of the activation markers CD137 and CD154 remained stable, except in CD4⁺ H3N2-TCPs, indicating preservation of antigen responsiveness during expansion (Figure 2B).

**Figure 2.**
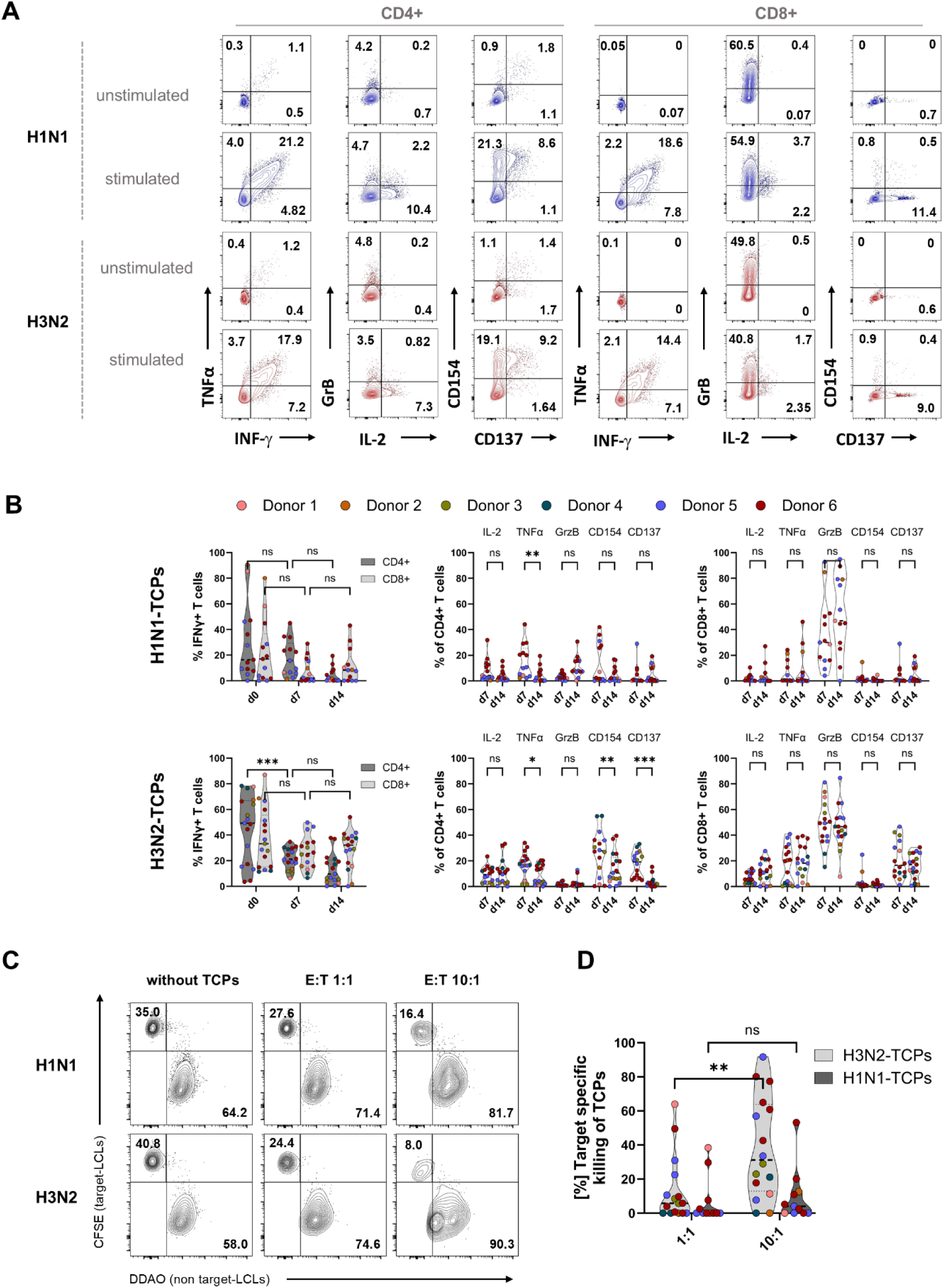
IAV-TCPs display effector functions *in vitro* and eliminate target-specific cells. **A)** Representative flow cytometry plots showing peptide stimulated H1N1-and H3N2-TCPs and unstimulated controls at d7 of expansion. Production of effector cytokines TNFα, IFN-γ, IL-2 and molecule granzyme B (GrzB) and expression of activation markers CD154 and CD137 are shown for CD4^+^ and CD8^+^ TCPs. **B)** Violin plots showing frequencies of effector cytokine production (IFN-γ, IL-2, TNFα), granzyme B and activation marker expression (CD154, CD137) for CD4^+^ and CD8^+^ populations relative to unstimulated controls at day 7 and 14. Upper panel: H1N1-TCPs (n=13, 4 donors); lower panel: H3N2-TCPs (n=17, 6 donors). Every dot per time point represents independent batch experiment. Donors are color coded according to the legend. Two-way ANOVA p<0.05 (*), p<0.01 (**), p<0.001 (***), p<0.0001 (****). **C)** Representative flow plots depicting gating strategy for killing assay. Gating was performed on living lymphocytes and T cells were excluded and gated only on lymphoblastoid cell lines (LCLs) as shown in Supplementary Figure 1B. Discrimination of target-LCLs (CFSE^+^) from non-target-LCLs (DDAO^+^) without or with TCPs at effector to target (E:T) ratio 1:1 and 10:1 for H1N1-and H3N2-TCPs. **D)** Quantification showing dose dependency of target-specific killing on d14 for H1N1-(n=11) and H3N2-(n=16) TCPs at E:T 1:1 and 10:1, respectively. Two-way ANOVA p<0.05 (*), p<0.01 (**), p<0.001 (***), p<0.0001 (****).

To evaluate the functional ability of IAV-TCPs to recognize and eliminate antigen-presenting target cells, a killing assay was performed at the endpoint of expansion (day 14) (Figure 2C and Supplementary Figure 1B). Consistent with the observed differences in phenotypic composition and cytokine production between the TCP subsets, H3N2-TCPs exhibited a higher cytotoxic capacity compared to H1N1-TCPs (Figure 2C,D). Both H3N2-and H1N1-TCPs demonstrated dose-dependent cytotoxic effects, with increased killing efficiency observed at higher E:T ratios (Figure 2D). Notably, at an E:T ratio of 10:1, H3N2-TCPs exhibited a mean target cell killing rate of 38%, whereas H1N1-TCPs displayed a lower killing efficiency <10%. These findings highlight the capacity of IAV-TCPs to mediate antigen-specific cytotoxicity, with distinct functional differences between H3N2-and H1N1-TCPs. The dose-dependent killing observed in both products underscore their potency, with the H3N2-TCPs emerging as the more efficient effector population, suitable to establish our 3D testing platform.

### Robustness of iPSC-derived alveolar lung organoid differentiation across different cell lines enables autologous 3D testing platform for virus infection

To generate human iPSC-aLOs, which serve as the model for our proof-of-concept platform, we assessed the reproducibility of differentiation across multiple human iPSC lines. We have applied a previously published protocol to the iPSC line BIHi289-B (autologous donor used in this study) and compared to the BU3NG line, as well as to several in-house reference iPSC lines (BIHi289-A, BIHi289-C, BIHi250-A, BIHi005-A, BIHi001-B, UCSFi001-A) ^40–42^.

Differentiations were monitored based on cell morphology and efficiencies assessed at day 3 (definitive endoderm) and day 14 (lung progenitor stage) by FACS (Supplementary Figure 2A,B). Lung progenitors displayed a high proportion of NKX2-1^+^/EpCAM^+^ cells, indicating the epithelial nature and overall efficiency of the differentiation. However, CPM expression varied between iPSC lines and across independent experiments (Supplementary Figure 2B). Despite this variability, iPSC-aLOs were successfully generated from all iPSC lines (Supplementary Figure 2C). iPSC-aLOs’ phenotype was stable upon passaging as shown by expression levels of pulmonary lineage markers (Supplementary Figure 2D).

Following the successful derivation of iPSC-aLOs, we evaluated their response to infection with human IAV strains H3N2 and H1N1 in comparison to well established adult stem cell derived alveolar-like lung organoids (ASC-aLOs) ^43^. Organoids were inverted into apical-out conformation, predominantly characterized by folded morphology, less defined lumen and tight junction ZO-1 localization change (Supplementary Figure 3A). Both organoid models were infected in parallel with both IAV strains H3N2 and H1N1, respectively. Viral replication was confirmed via a plaque assay showing that replication dynamics of the iPSC-aLOs were comparable with those of the ASC-aLOs for the respective IAV strains (Supplementary Figure 3B). As in the previous study ^43^, viral titers at 16 hours post infection (hpi) varied between ∼10^6^ PFU/ml for H3N2 and ∼10^5^-10^6^ PFU/ml for H1N1 for both organoid models. To further investigate the infection response of iPSC-aLOs, we performed single-cell RNA sequencing (scRNA-seq) using uninfected and H3N2-infected iPSC-aLOs at 24hpi (Figure 3A). H3N2-infection was confirmed by plaque assay for the investigated lines BU3NG and BIHi289-B (Figure 3B).

**Figure 3.**
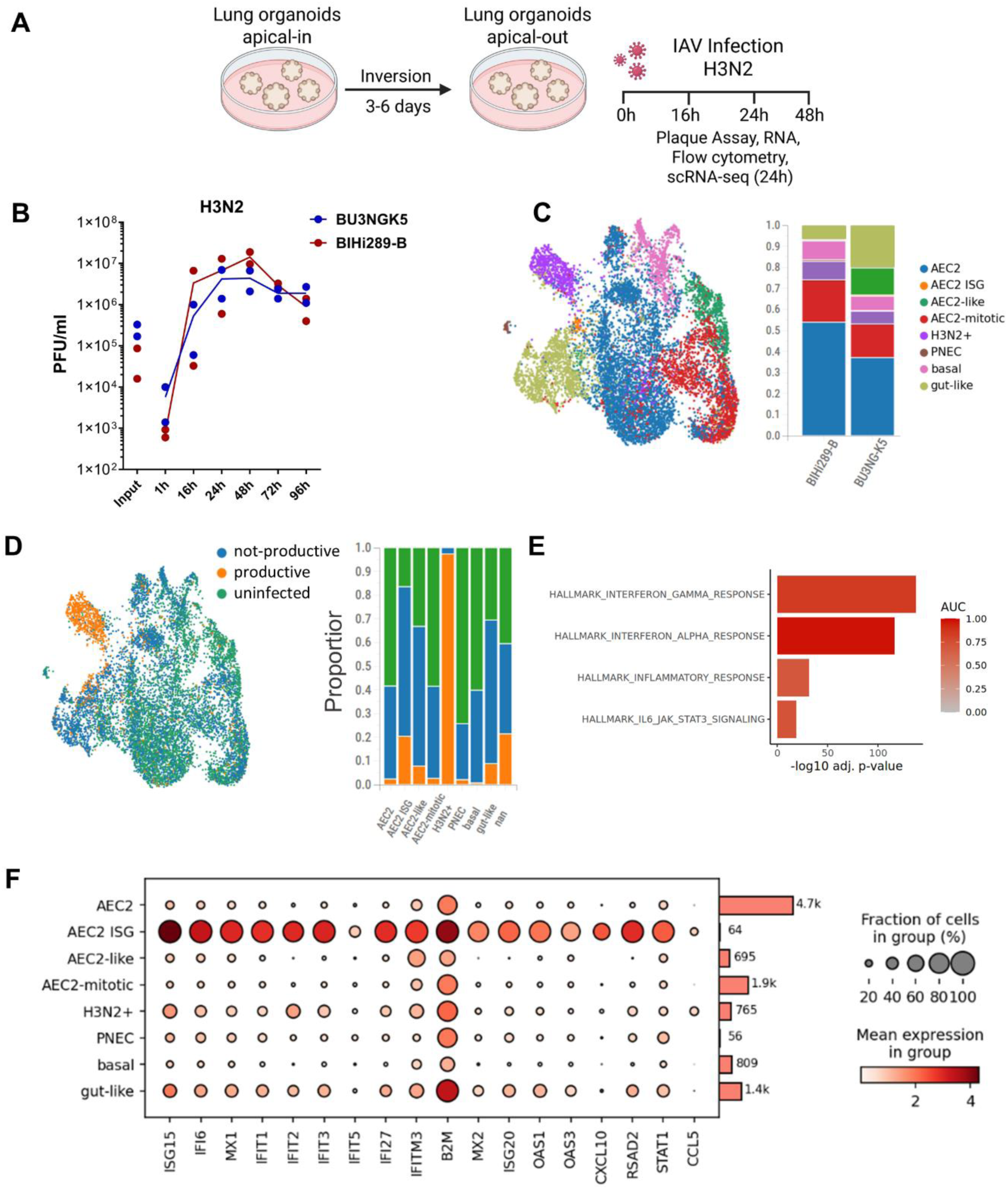
**H3N2-infected iPSC-aLOs show an increased IFN response**. **A)** Experimental design scheme for apical-out inversion of iPSC-aLOs and endpoint assays upon H3N2 infection. The image was created with BioRender.com. **B)** Quantification of Plaque Forming Units (PFU) showing kinetic of H3N2 replication in iPSC-aLOs derived from two different hiPSC lines. Mean values for each group are depicted by lines with dots for individual donors at indicated timepoints (n=2 independent experiments per hiPSC line). **C)** UMAP showing iPSC-aLOs composition (left) and stacked bar plots depicting cell population frequencies across iPSC-aLOs derived from two hiPSC lines (right) after single-cell RNA sequencing (scRNA-seq) at 24 hours post infection (hpi). **D)** UMAP derived from scRNA-seq showing not productive or early infected, likely productive and uninfected cells distribution (left) and stacked bar plots showing their frequencies across cell types present in iPSC-aLOs (right). **E)** Gene set enrichment analysis (GSEA) displaying correlation of differential gene expression analyzed by scRNA-seq between H3N2-infected and non-infected iPSC-Los 24hpi and IFN**-**γ, INF-α, inflammatory response and IL6/JAK/STAT3 signaling signatures (AUC= area under ROC curve, being 1 highest correlation). **F)** Dot plot derived from sc-RNAseq analysis showing expression of IFN stimulated genes per cell type cluster.

Cluster annotation was performed using marker genes from the literature, confirming that the predominant cell population within the iPSC-aLOs consists of alveolar-like epithelial type II cells (AEC2), with different subtypes including AEC2-like and AEC2-mitotic progenitors (Figure 3C and Supplementary Figure 3C,D) ^41,43^. Interestingly, we could determine a segregated cluster of AEC2 showing up-regulation of interferon stimulated genes (AEC2-ISG high). In addition, cells belonging to the H3N2+ cluster were highly infected and could not be assigned to a specific cell type since viral transcripts were dominant as marker genes. In addition to AEC2 populations, the analysis also identified pulmonary neuroendocrine cells (PNECs), basal cells and gut-like cells, suggesting some degree of lineage diversification (Figure 3C and Supplementary Figure 3C,D). The distribution of identified cell populations was consistent among iPSC-aLOs derived from the two iPSC lines (Figure 3C). Apical-in uninfected controls included in the analysis were compared with the apical-out condition to confirm both conformations kept similar cell type composition and cell cycle state (Supplementary Figure 3E).

Viral transcripts were detected across all cell populations (Supplementary Figure 4A). Similar to previous reports, we used the simultaneous co-expression of four key IAV genes (PA, PB1, PB2, and NP) within individual host cells, to differentiate a likely productive from a likely not productive infection ^43,44^. Likely productive infected cells are expected to co-express all four viral genes at detectable levels, while cells expressing only a subset of these genes were considered potentially to be in early stages of viral replication. All populations contained both likely productive and not productively infected cells. The AEC2-ISG high cluster was composed mostly of not productively (early stage) infected cells and the H3N2⁺ cluster was predominantly composed of likely productively infected cells (Figure 3D, Supplementary Figure 4B). Differential gene expression between infected and uninfected samples showed a distinct immunological interferon (IFN) response as shown by hallmark pathway analysis (Figure 3E, Supplementary Figure 4C, Supplementary Table 1). The IFN response signature was greater in AEC2-ISG^high^ and H3N2^+^ clusters with similar levels in likely productively and not productively infected cells (Figure 3F, Supplementary Figure 4D). The iPSC-aLOs’ IFN response based on selected ISGs (*IFIT1, IFIT3, IFI6, ISG15* and *MX1)* was comparable to the one of ASC-aLOs, following viral challenge with either H3N2 or H1N1, showing similar suitability to model IAV infections (Supplementary Figure 5). For iPSC-aLOs, presentation of antigens and forming immunological synapses with T cells, HLA molecules expression is of major importance. Consistent with their epithelial origin, iPSC-aLOs showed low HLA class II expression suggesting limited capacity for MHC class II–mediated antigen presentation (Supplementary Figure 6A,B). However, like in ASC-aLOs, robust HLA class I expression was observed in iPSC-aLOs indicating presentation capacity of intracellular viral antigens to CD8^+^ T cells. Thus, iPSC-aLOs effectively model viral replication and IFN responses and are well-suited for studying cytotoxic T cell-mediated immunity in a physiologically relevant *in vitro* system.

### H3N2-infected 3D organoid co-culture system: proof-of-concept approach for efficacy and safety assessment of antiviral TCP-mediated cytotoxicity

To establish a co-culture system for studying IAV-TCP-mediated killing in a controlled and almost matrix free environment compatible with automated imaging, we optimized our organoid culture conditions by transitioning to a 96-well plate format containing 120 microwells per well (Gri3D) (Figure 4A). Due to the low cell numbers and small well volumes in the Gri3D format, we first conducted proof-of-concept experiments to evaluate its suitability for infection studies and to optimize infection conditions and viral sampling. ASC-aLOs were chosen for their uniform morphology in Gri3D and higher number of cells per organoid (Supplementary Figure 7A). We compared continuous supernatant sampling from the same well over time with single-timepoint sampling per well. Plaque assays confirmed successful infection and viral detection using both methods (Supplementary Figure 7B). To reduce well-to-well variability, the continuous sampling approach was used in subsequent experiments. Media formulations supporting both IAV-TCPs and iPSC-aLOs maintenance were tested. For this, TCPs functionality was prioritized. Hence, TCPs were transitioned into various test media and evaluated for viability and functionality (Supplementary Figure 7C). Based on the optimal balance, a 1:1 mixture of iPSC-aLO basal medium (cSFDM) and T cell medium (TCM) was selected for all co-cultures. Effector to target ratios were tested by titration of T cell numbers at seeding for better optical resolution per iPSC-aLO (Supplementary Figure 7D). Seeding of 50,000 T cells per well of Gri3D was selected for subsequent experiments.

**Figure 4:**
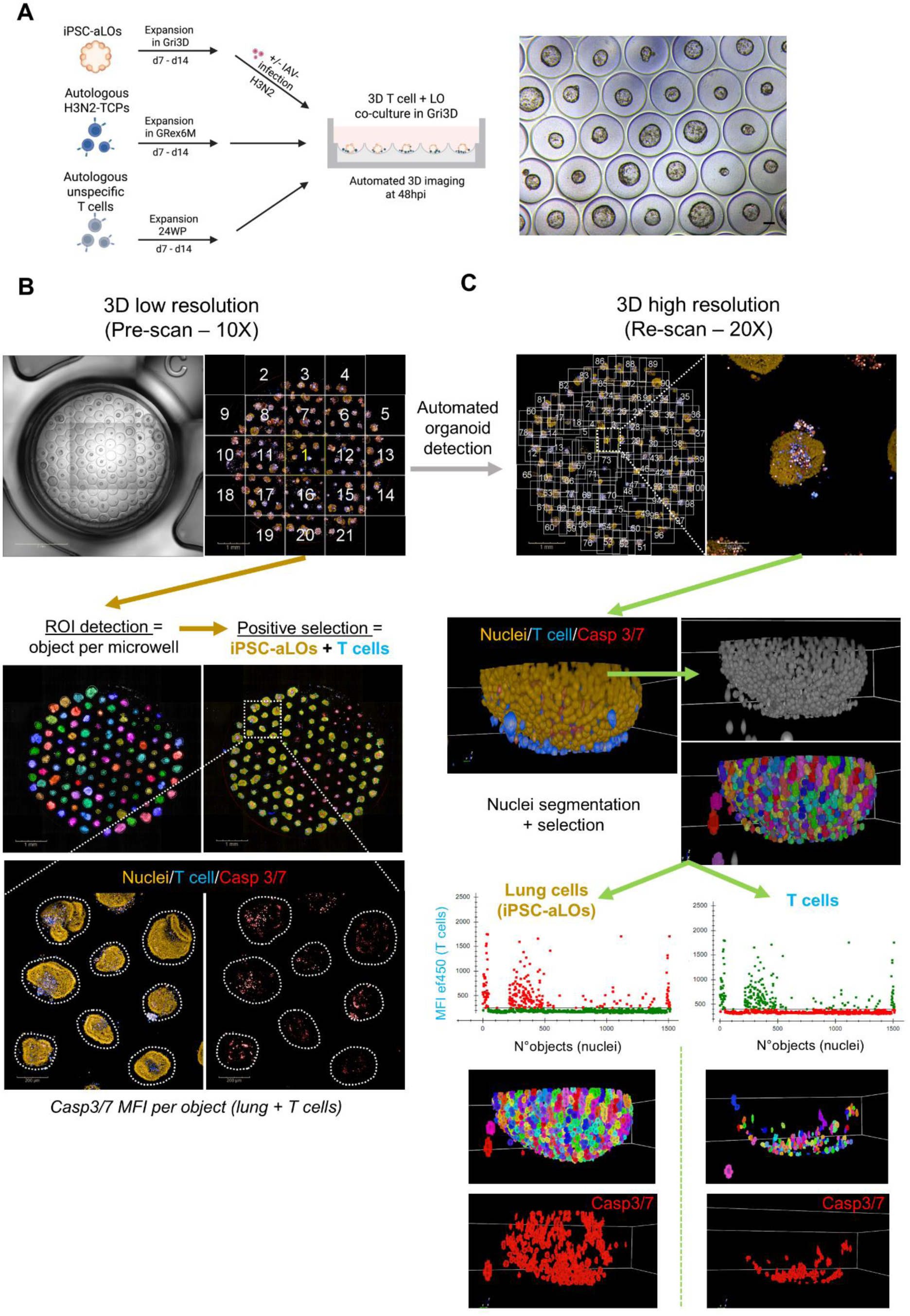
P**r**inciple **of automated 3D assay for safety and efficacy assessment of TCPs. A)** Experimental design scheme showing co-culture strategy using microwells (Gri3D) for infection and read-out by high-throughput imaging and high-content analysis. The image was created with BioRender.com (left). Example of bright field image showing iPSC-aLOs (at day 9 of expansion in Gri3D) prior infection, scale bar: 100µm (right). **B)** Representative picture showing whole well lower resolution pre-scan and downstream analysis example to determine Caspase 3/7 mean fluorescence intensity per region of interest (ROI) composed by iPSC-aLO + T cells. Scale bars: 1mm and 200µm. **C)** Pre-scan from panel B used to localize XYZ coordinates for re-scan at higher magnification followed by analysis based on nuclei segmentation and recognition. Caspase 3/7 signal at higher resolution can be assigned to iPSC-aLO cells or T cells.

To assess autologous H3N2-TCP functionality in our 3D system, we developed an imaging-based assay involving a low-magnification pre-scan to locate all organoids, followed by higher-magnification XYZ re-scan acquisition of individual organoids (Figure 4B,C). For analysis, we systematically compared low-versus high-resolution image data. Low-resolution analysis maximum projections were used to define regions of interest (ROIs) containing one iPSC-aLO and T cells for which caspase 3/7 median fluorescence intensities (MFIs) were calculated (Figure 4B). High-resolution 3D image stacks enabled volumetric nuclear segmentation and discrimination between T cells and iPSC-aLO-derived cells using fluorescent thresholds (Figure 4C). Caspase 3/7⁺ cell frequencies could then be quantified for each individual population (Figure 4C).

The newly developed analysis pipelines at low and high resolution were used to assess H3N2-TCP-mediated cytotoxicity on iPSC-aLOs (BIHi289-B) in an autologous set up (Figure 4A). In the low-resolution analysis, uninfected iPSC-aLOs co-cultured with unspecific T cells showed a significant increase in caspase 3/7 activity compared to iPSC-aLOs alone, with a further increase upon co-culture with H3N2-TCPs (Figure 5A). In infected samples, caspase 3/7 levels rose further, reaching the highest levels in the presence of H3N2-TCPs. These patterns were observed at both MOI 1 and MOI 10, with a marked increase at the higher MOI, indicating a dose-dependent effect. Maximum projection in low resolution imaging analysis did not allow discrimination of apoptotic signals from iPSC-aLOs or T cells (Figure 5A). High-resolution analysis confirmed the overall trends observed in the low-resolution data, particularly the marked increase in caspase 3/7 activity in infected iPSC-aLOs co-cultured with H3N2-TCPs (Figure 5B,C). However, it revealed more distinct differences in the uninfected controls. Specifically, caspase 3/7⁺ lung cells remained lower in uninfected iPSC-aLOs, with only modest increases upon co-culture with unspecific T cells or H3N2-TCPs, showing most of the apoptotic signal detected at low magnification belonged to T cells (Figure 5C). In infected samples, both unspecific T cells and H3N2-TCPs contributed to increased apoptosis, with H3N2-TCPs consistently inducing the highest caspase 3/7 activity across both MOIs and analysis methods (Figure 5 A,B,C). The low-resolution analysis could not differentiate between iPSC-aLO-derived cells and T cells within the ROI, limiting cell-type-specific interpretation (Figure 5A). In the high-resolution analysis, caspase 3/7 quantification within the T cell populations revealed generally elevated apoptosis levels. H3N2-TCPs exhibited a significantly higher mean frequency of caspase 3/7⁺ cells (50-60%) compared to unspecific T cells (20-40%) (Figure 5C). These findings highlight the added value of single-cell resolution in accurately assessing cell-type-specific responses within complex co-culture systems.

**Figure 5.**
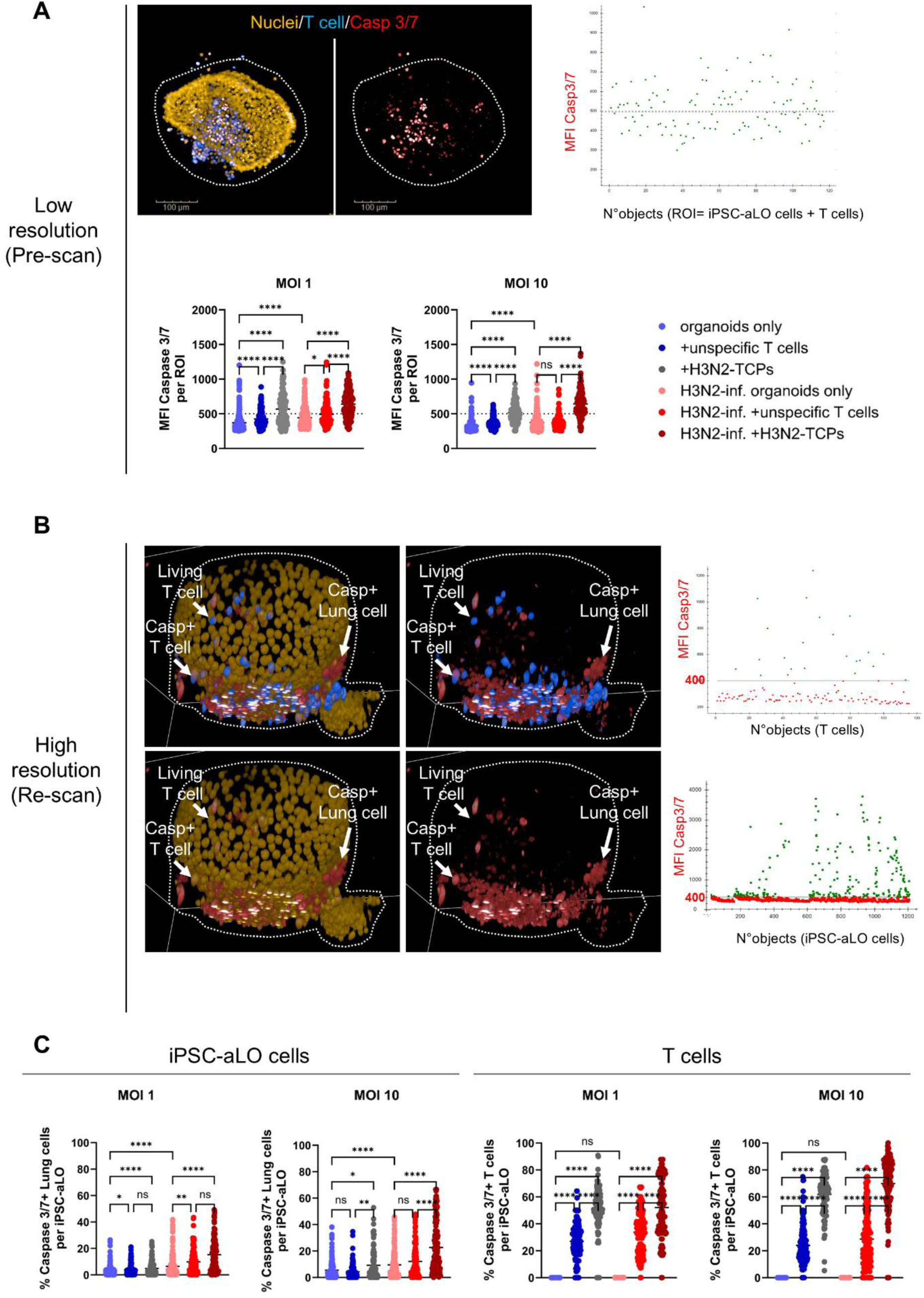
H3N2-infected 3D organoid co-culture system shows efficacy of antiviral H3N2-TCPs. **A)** Representative image of one region of interest (iPSC-aLO + T cells) showing Caspase 3/7 (upper left) and scatter plot for Caspase 3/7 mean fluorescence intensity (MFI) from H3N2-infected iPSC-aLO + H3N2-TCP (upper right). Plots showing Caspase 3/7 MFI quantification per ROI for MOI=1 and 10. Dots represent technical replicates of individual organoids for two independent experiments (n=2). Two-way ANOVA Kruskal-Wallis Test p<0.05 (*), p<0.01 (**), p<0.001 (***), p<0.0001 (****). Scale bars: 100µm. Dotted line serves as reference between scatter plot and quantification plots. **B)** Representative 3D image of one iPSC-aLO and T cells in co-culture showing Caspase 3/7 at single cell level (left) and scatter plots for Caspase 3/7 MFI (cut-off= 400) per cell type, respectively (left). **C)** Plots showing percentage of Caspase 3/7 positive cells in lung cells (iPSC-aLO cells) and T cells for MOI=1 and 10. Dots representing technical replicates of individual organoids for two independent experiments (n=2). Two-way ANOVA Kruskal-Wallis Test p<0.05 (*), p<0.01 (**), p<0.001 (***), p<0.0001 (****).

## Discussion

In this study, we established a GMP-compliant workflow for generating IAV-TCPs targeting IAV subtypes H3N2 and H1N1 and performed comprehensive phenotypic and functional characterization. In parallel, we derived iPSC-aLOs, validated their infection susceptibility, and applied them in an autologous 3D co-culture system. This platform enabled us to assess TCP-mediated cytotoxicity in a physiologically relevant context. Together, our findings demonstrate core functionalities of IAV-TCPs *in vitro*, the antiviral responsiveness of IAV-infected iPSC-aLOs and provide evidence for target-specific elimination of infected cells by H3N2-TCPs in a 3D human alveolar-like infection model.

We present an IAV-TCP manufacturing approach, which despite challenges such as the limited abundance of the target T cell populations in peripheral blood with low initial precursor frequencies, remains feasible due to advancements in cultivation techniques, such as the G-Rex6M system ^45,46^. The platform offers key advantages for GMP-compliant TCP manufacturing, particularly by reducing the need for frequent culture manipulation due to the large media volumes per vessel. This minimizes external culture interference, promotes process standardization and supports the scalable expansion of IAV-TCPs. Phenotypic analysis of the product revealed a CD4-dominated profile, in contrast to previous TCPs specific to CMV-or SARS-CoV-2, expanded under similar media conditions ^11,32–34^. However, since our approach primarily utilized peptides spanning conserved proteins (NP, MP1, MP2), our observed findings align with prior studies, demonstrating that responses to these immunodominant epitopes are predominantly mediated by CD4^+^ T cells ^37,47–50^. Our functional assays confirmed that IAV-TCPs retained their effector functions throughout expansion, as demonstrated by cytokine production and upregulation of activation markers. Overall, frequencies either remained stable or decreased slightly, supporting their potential for downstream applications in adoptive immunotherapy. Although the CD4-dominated phenotype initially raised concerns regarding cytotoxic potential of these products, both IAV-TCPs demonstrated target-specific killing in a suspension culture assay, exhibiting dose-dependent cytotoxicity. These findings are consistent with prior studies, reporting that CD4^+^ T cells can mediate cytotoxicity in IAV infection ^37,48,51^. Notably, IAV-TCP killing efficiencies were lower compared to CMV-or SARS-CoV-2-specific TCPs ^32–34^. We speculate that this difference stems from the antigenic characteristic of IAV evolution, which drives immune selection for highly cross-reactive T cells that predominantly support antibody responses against viral surface proteins ^37,52,53^. Further research is needed to identify the most immunodominant epitopes, correlate these with MHC-T cell subtype interactions, and assess their impact on viral clearance and immune response efficacy. Establishing these correlations will not only enhance IAV-TCP manufacturing and epitope selection but provide deeper insights for the design and development of novel flu vaccines and other preventive therapies.

Robust generation, comprehensive characterization and H3N2 susceptibility of iPSC-aLOs demonstrated their suitability as a relevant host model, promoting similar viral replication and IFN responses to ASC-aLOs, despite their early developmental stage ^43,54^. Strong MHC class I but low MHC class II expression observed in iPSC-aLOs, supports modeling CD8^+^ T cell responses but limits CD4^+^ T cell activation. Despite low CD8^+^ T cell frequencies in TCPs, iPSC-aLOs’ MHC I profile enabled investigation of cytotoxic T cell activity. Future integration of immune cells, such as alveolar macrophages, may overcome this in the future ^55–57^.

Nearly matrix-free co-culture of organoids with T cells in Gri3D facilitated standardized organoid size distribution, prevented fusion and minimized movement, ensuring more defined and close interactions between infected organoids and H3N2-TCPs, while maintaining reproducibility across experiments enabling automated high throughput high content imaging. Since the natural route of IAV infection occurs at the apical surface of alveolar epithelial cells, in extracellular embedded set up organoids were inverted into apical-out conformation to facilitate successful infection, step which is no longer needed in Gri3D ^43,44,58^. However, the system posed challenges to detect viral replication. Plaque assays performed from supernatants using continuous sampling was possible for ASC-aLOs but not for iPSC-aLOs cultured in Gri3D, likely due to the lower cell density of iPSC-aLOs causing viral titers to fall below the detection limit of the plaque assay. A more sensitive approach, such as PCR-based detection, may allow reliable viral quantification. Alternatively, the use of fluorescently tagged viruses for imaging would leverage the system by being able to follow infected cells along the process ^59,60^. Despite the limitation to reliably quantify viral replication in iPSC-aLOs cultured in Gri3D, our imaging-based analysis provided strong evidence of H3N2 infection of iPSC-aLOs, showing dose dependent increase in caspase 3/7^+^ cells upon interaction with H3N2-TCP.

For image analysis, we systematically compared low-(pre-scan) and high-(re-scan) resolution acquisitions to assess their predictive accuracy for evaluating TCP safety and efficacy. Both analysis methods effectively identified general trends, including increased apoptosis in infected conditions and the relative safety of the product when compared to unspecific T cell controls. Lower-resolution analysis proved valuable in capturing broad patterns of TCP efficacy at approximately 5X higher acquisition speed than high-resolution, however, it lacked the precision required for accurately assessing safety, primarily due to its inability to distinguish between lung epithelial cells from iPSC-aLOs and T cells within the ROIs. As T cells adjacent to iPSC-LOs could not be excluded from analysis, this limited the interpretation of cell-type-specific apoptosis. In contrast, high-resolution analysis enabled more precise quantification of apoptosis within distinct cell populations, offering deeper insights into both the frequency and distribution of caspase 3/7⁺ cells. This included a clearer understanding of T cell-specific activity, with H3N2-TCPs showing a significantly higher apoptotic effect on infected lung cells compared to unspecific T cells, showing that H3N2-TCPs likely execute their effector functions in a target-specific manner. These findings emphasize the importance of higher imaging resolution for accurate cell-type discrimination and robust evaluation of product specificity. Moreover, high-resolution data raised additional questions, such as the mechanism behind the increased apoptosis observed in infected iPSC-aLOs co-cultured with autologous unspecific T cells compared to infected iPSC-aLOs alone. One possible explanation is TCR-independent pre-activation of unspecific T cells, potentially induced during expansion. Alternatively, factors secreted by infected iPSC-aLOs may have contributed to bystander activation ^61^. MOI increment from 1 to 10 amplified H3N2-TCP-mediated apoptosis, indicating a dose-dependent relationship between infection burden and cytotoxic response. However, further refinement of the assay is required to enable controlled application of defined E:T ratios, which would allow for a more precise assessment of TCP dose-responsiveness. To comprehensively address safety concerns, a more detailed investigation of both on-and off-target effects is required. As a next step, we propose the use of a fluorescently labeled IAV strain to definitively confirm infection status and enable clear visual distinction between infected and uninfected lung cells ^59,60^. This refinement would significantly enhance the assay’s predictive value by allowing more accurate correlation of T cell activity with viral clearance. Ideally, the assay should incorporate live cell imaging to provide dynamic insights into the kinetics and pharmacodynamics of TCPs. However, implementing live cell imaging presents technical challenges, particularly due to the prolonged scanning time required for high-resolution imaging of multiple 3D organoids. As a result, live cell imaging would currently be feasible only for a limited number of organoids, limiting scalability. In future experiments, testing TCPs with different specificities, such as H1N1-specific T cells, will be essential to further validate product specificity and assess potential cross-reactivity. Comprehensive T cell profiling following co-culture, including analysis of activation markers, cytokine production, and exhaustion status via flow cytometry, will yield critical insights into TCP functionality and persistence.

Overall, our proof-of-concept findings show for the first time the feasibility of integrating antiviral TCPs with lung organoid infection models to assess immune-mediated cytotoxicity in a 3D context. This approach offers a promising platform for studying antiviral immunity, evaluating candidate therapies, and advancing insights into host-pathogen interactions within a physiologically relevant *in vitro* system. We propose this assay as a complementary follow-up to conventional suspension-based killing assays, particularly for validating TCP candidates that demonstrate efficacy in initial screens. Our platform provides a foundation for deeper investigation into TCPs mechanisms of action, safety, and efficacy.

## Methods

### Blood donors and samples

The study was approved under permit EA1/052/22 and all donors provided written informed consent. Peripheral blood mononuclear cells (PBMCs) were isolated from heparinized venous blood of healthy donors using Pancoll Cell-Separating Solution (PAN-Biotech) and density gradient centrifugation following manufacturer indications.

### Manufacturing and expansion of IAV-TCPs

IAV-TCPs were generated using an adapted version of previously described protocols ^11,32–34^. In brief, PBMCs were isolated from 50 ml of heparinized venous blood by Pancoll density gradient centrifugation (PAN-Biotech) and seeded into complete medium VLE RPMI 1640 (PAN-Biotech) supplemented with 100 IU/mL penicillin and 100 µg/ml streptomycin (Gibco), 10% fetal calf serum, 10 ng/mL recombinant human IL-7 (rhIL-7, CellGenix), and 10 ng/mL recombinant human IL-15 (rhIL-15, CellGenix) in 24-well plates. PBMCs were stimulated for 6 hours with 0.5 µg/mL overlapping peptide pools of individual IAV proteins (JPT Peptide Technologies), including MP1, MP2, NP, and HA for H1N1, or MP1, NP, and HA for H3N2. Interferon-γ (IFN-γ) secreting cells were subsequently labeled and isolated using the IFN-γ secretion assay (Miltenyi Biotech) according to the manufacturer’s instructions. Autologous feeder cells were generated from the respective IFN-γ^-^ cell fraction by irradiation (30 Gy using a GSR D1 (Gamma-Service Medical GmbH)). IFN-γ^+^ cells including 25×10^6^ feeder cells were cultivated in expansion medium composed of 1:1 Advanced RPMI 1640 Medium (Gibco) and Click’s Medium (Fujifilm) supplemented with 1% GlutaMAX (Gibco), with 100 IU/ml penicillin and 100 µg/ml streptomycin (Gibco) and 10% fetal calf serum (FCS, PAA), 10 ng/ml recombinant human (rh) IL-7 and rhIL-15 (CellGenix) in G-Rex6M Well Plates (ScaleReady) and cultivated in humidified incubators at 37°C and 5% CO_2_. Medium changes and cell passaging were performed every 7 days. Glucose and lactate concentrations were measured every 2 days using a Lactate Plus device with strips (Nova Biomedical) and a Glucose device with strips (Bayer) to monitor culture stability during expansion.

### Lymphoblastoid Cell Line Generation

Autologous lymphoblastoid cell lines (LCLs) from each donor were generated by infecting 5×10^6^ PBMC with concentrated virion containing supernatant of the cell line producing EBV B95.8, as previously reported ^11,32–34,62^. Infection was performed in complete medium supplemented with 0.5 μg/ml cyclosporin A and 4 μg/ml CpG in a 24-well plate. Stably growing LCLs were continuously cultured in complete medium in 75cm^3^ cell culture flasks. Half of the medium was replaced once to twice a week with fresh medium.

### Phenotypic and functional flow cytometric assessment of IAV-TCPs

For assessment of the IAV-TCPs phenotype, cytokine production and activation, cells were collected in complete medium without cytokines and rested overnight in 5ml round bottom centrifugation tubes. For stimulation, 1×10^4^ LCLs and 0.5 µg/mL of the respective overlapping peptide pools were added to 1×10^6^ TCPs. LCLs and respective volumes of DMSO were added to unstimulated controls. Intracellular cytokine production was captured by addition of 2 mg/mL of Brefeldin A (Sigma-Aldrich) after 1h of stimulation. Extra-and intracellular antibody staining was performed with all antibodies from BioLegend, unless stated otherwise. Surface antibody staining was performed after 6h of stimulation with human anti-CCR7 (G043H7),-CD45RA (HI100) and LIVE/DEAD Fixable Blue Dead Cell Stain (L/D; Invitrogen). According to the manufacturer instructions, cells were fixed and permeabilized using the FoxP3/Transcription Factor Staining Buffer Set (eBioscience) and subsequently stained intracellularly with human anti-CD3 (OKT3),-CD4 (SK3),-CD8 (SK1), - IFN-γ (4S.B3, eBioscience, 00-5523-00),-TNF-α (MAb11),-IL-2 (MQ1-17H12),-CD137 (4-1BB),-CD154 (24-31) all from BioLegend.

### Killing assay

Target specific killing was determined by flow cytometric killing assay as previously described^34^. Briefly, autologous target LCLs were stained with 10 mM carboxyfluorescein diacetate succinimidyl ester (CFSE-DA; Sigma-Aldrich) for 4min, whereas non-target LCLs were stained with 5 mM Cell Trace Far Red (DDAO, Invitrogen) for 10min. Target LCLs were loaded for 2h with 2 µg/mL of the respective overlapping peptide pools and equally mixed with non-target LCLs. In complete medium rested TCPs were co-cultivated at distinct ratios with LCL mixes. T cell-free LCL mixtures served as internal controls. After 16h, co-cultures were stained with L/D to exclude dead cells from the analysis. All flow cytometry samples were analyzed using a CytoFLEX flow cytometer (Beckman Coulter) and FlowJo-10 software v10.8 (Tree Star).

### iPSC lines, culture and maintenance

The BU3 human iPSC line carrying the NKX2-1^GFP^ reporter (BU3NG), was generously gifted by the Kotton Lab (Boston University) ^40,41,63^. All iPSC lines used in this study (BU3NG, BIHi289-A; https://hpscreg.eu/cell-line/BIHi289-A, BIHi289-B; https://hpscreg.eu/cell-line/BIHi289-B, BIHi289-C; https://hpscreg.eu/cell-line/BIHi289-C, BIHi250-A; https://hpscreg.eu/cell-line/BIHi250-A, BIHi005-A; https://hpscreg.eu/cell-line/BIHi005-A, BIHi001-B; https://hpscreg.eu/cell-line/BIHi001-B, UCSFi001-A; https://hpscreg.eu/cell-line/UCSFi001-A) belonged to quality-controlled banks and displayed a normal karyotype. iPSC lines were maintained in feeder-free conditions on Geltrex (Gibco, 1:120) coated 6-well plates in either Essential 8 medium (E8) or mTeSR1 (Stem Cell Technologies), passaged as clumps using 0.5 mM EDTA (Invitrogen) every 3-4 days and cultivated at 37°C, 5% CO_2_, 5% O_2_ in humidified incubators ^64,65^. Before differentiation, iPSCs were adapted for 1-2 passages to mTeSR1 media if they came from E8. Colonies were cultivated up to a confluency of 70-80% to ensure the cells have reached appropriate density and are ready to use for differentiation.

### Differentiation of human iPSCs into NKX2-1+ lung progenitors

Human iPSC lines (BU3NG, BIHi289-A, BIHi289-B, BIHi289-C, BIHi250-A, BIHi005-A, BIHi001-B, UCSFi001-A) were differentiated into NKX2-1^+^ lung progenitor cells as reported ^41^. All bright field images from culture monitoring were acquired using Leica DMi8 microscope.

### MACS sorting of CPM^+^ lung progenitor cells

Between days 14-16 of differentiation, cells were collected for MACS sorting of NKX2-1^+^ lung progenitors using anti-human CPM (FUJIFILM). Cells were detached using 0,25% Trypsin/EDTA 2 mM (Gibco), facilitated by scratching the culture in a cross shape and incubated for 5-7min at 37°C. Single cell suspension was collected and washed in FACS buffer. Cells were gently re-suspended and passed through a 40µm cell strainer. Single cells were surface stained with anti-human CPM (1:100) for 45min at 4°C, washed and re-suspended in 80µl FACS buffer per up to 10^7^ cells. CPM was captured using 20µl Anti-Mouse IgG2a+b Microbeads (Miltenyi Biotec) per up to 10^7^ cells and incubated for 15min at 4°C. Separation was performed using MS separation columns. Recovered cells were replated in growth factor-reduced Matrigel (“GFR Matrigel”; Corning) at 50,000-150,000 CPM+ cells per 50 µl droplet in 24-well plates, depending on the line. Encapsulated CPM^+^ cells were covered with alveolar differentiation medium, “CK+DCI” containing a base of complete serum free differentiation media (cSFDM)^41^ with 3 µM CHIR99021 (Biovision or Abcam), 10 ng/m rhKGF (R&D Systems), 50 nM dexamethasone (Sigma-Aldrich), 0.1 mM 8-bromoadenosine 3′,5′ cyclic monophosphate sodium salt (Sigma-Aldrich), 0.1 mM 3-isobutyl-1-methylxanthine (IBMX, Sigma-Aldrich), supplemented with 10 µM Y-27632 (Rho-kinase inhibitor (ROCKi) for 72h post sort. Media change using CK+DCI media was performed every 48-72h. Sorting quality assessment was performed by flow cytometry. Cells were surface stained with anti-human EpCam (REA764, Miltenyi Biotec), then fixed and permeabilized using the FoxP3 Staining Buffer Kit (Miltenyi Biotec) and subsequently stained intracellularly with anti-human NKX2-1 antibody (EP1584Y, Abcam) for 45min at 4°C. Secondary labelling was performed using donkey anti-mouse Alexa Fluor 488 (Thermo Fisher) and donkey anti-rabbit Alexa Fluor 647 (Thermo Fisher). Samples were analyzed using a CytoFLEX flow cytometer (Beckman Coulter) and FlowJo-10 software v10.8 (Tree Star).

### Long term culture, passaging and freezing of iPSC-aLOs

Human iPSC-aLOs developed within 1-2 weeks after CPM+ cells encapsulation into Matrigel domes and were passaged and expanded in CK+DCI medium for weeks up to months. Medium was refreshed every 48-72h and serial passaging performed every 10-14 days. For Passaging, Matrigel domes were dissociated for 30min at 37°C using Dispase 2 mg/ml (Roche). Retrieved organoids were either dissociated mechanically in clumps or enzymatically into single cell using 0,25% Trypsin/EDTA for 10min at 37°C. Single cells were re-seeded at 50.000-150.000 cells per 50 µl dome. Clumps were re-plated at different ratios (1:2-1:8) according to initial density. The medium was supplemented with 10 µM ROCKi for the first 48-72h after every passaging. For freezing, iPSC-aLOs were gently mechanically dissociated and frozen in FBS and 10% DMSO. Samples were collected over passages using lysis buffer from Aurum RNA Mini Kit (BioRad) and kept at-80°C until use. All bright field images from culture monitoring were acquired using Leica DMi8 microscope.

### ASC-aLO generation

Tumor-free peripheral lung tissue was obtained from patients undergoing lung resection surgery at collaborating hospitals in Berlin, Germany. The study was approved by the ethics committee of Charité-Universitätsmedizin Berlin (project EA2/079/13), and written informed consent was obtained from all participants. Human lung organoids were established as previously published ^43^. In short, tissue was washed, minced, and digested in 500 U/ml Collagenase type I (Gibco), 5 U/ml Dispase II (Gibco), and 1 U/ml DNase I (AppliChem) in HBSS^+/+^ containing 10 µM ROCKi (Tocris) for 60–90min at 37°C in a shaking water bath. Following filtration and red blood cell lysis using RBC lysis buffer (Invitrogen), single-cell suspensions were prepared for fluorescence-activated cell sorting.

HTII-280⁺/EpCAM^+^ alveolar epithelial cells were labeled using anti-HTII-280 IgM (Terrace Biotech, TB-27AHT2280), anti-EpCAM-PE (BioLegend, 324206) and secondary goat anti-mouse IgG (H+L) Alexa Fluor 488 antibody (Thermo Fisher, A-21042). Cells were sorted at FCCF DRFZ Core facility using FACS Aria II (BD Biosciences) and embedded in Cultrex (R&D Systems) at ∼1000cells/μl. After gelation at 37°C for 25min, organoid cultures were overlaid with basal medium (Advanced DMEM/F12, 10 mM HEPES, GlutaMax]) supplemented with 1xB27, 1xN2 (all Thermo Fischer), 10% R-spondin1 conditioned medium, 1.25 mM N-acetylcysteine (Sigma), 5 mM nicotinamide (Sigma), 0.5 μM SB202190 (Sigma), 1 μM A83-01 (Merck), 100 ng/ml Noggin (Gibco), 100 ng/ml FGF10 (Gibco), 25 ng/ml FGF7 (Gibco), and 3 μM CHIR99021 (Sigma). ROCKi (10 μM, Tocris) was included during the first three days of culture only. Organoids were maintained at 37°C, 5% CO₂, and passaged every 2–4 weeks by enzymatic dissociation using TrypLE Express (Gibco).

### RT-qPCR

RNA extracted using Aurum RNA Mini Kit (BioRad) followed by DNAse treatment using TURBO™ DNase (Invitrogen). cDNA was generated from 600 ng RNA per sample by reverse transcription using iScript cDNA Synthesis Kit (BioRad) according to the manufacturer’s instruction. Diluted cDNA (1:15) was used for qPCR, technical replicates of 4µl reactions were used for targeted amplification using SYBRGreen PCR Master Mix (Thermo Fisher). Primers sequences are described in Supplementary Table 2. RT-qPCR was run in Quant Studio 6 (Thermo Fisher). Relative gene expression was calculated using delta-delta Ct method and normalized to control sample.as described in figures for each experiment type.

### Virus strains

Human seasonal influenza virus A (IAV) A/Panama/2007/1999 (Pan/99[H3N2]) and A/Puerto Rico/8/34 (PR8 NS1-GFP[H1N1]) were kindly provided by Dr. Thorsten Wolff (RKI, Berlin) and propagated on MDCKII cells as described ^66^. Viral stocks were aliquoted, stored at −80°C, and titrated on MDCKII cells by plaque assay.

### Infection of lung organoids (LOs)

For infection, LOs (iPSC-aLOs or ASC-aLOs) were switched to apical-out (AO) conformation. Briefly, LOs were collected on ice using Corning Recovery solution (Corning), disrupted by repeated resuspension and incubated for 30min on ice on a shaking rotator. Recovered LOs were collected, washed with PBS-/-(Gibco) and cultured in suspension for 3-5 days in their respective medium, supplemented with 0.015% Gellan gum (VWR) on Anti-Adherence Rinsing Solution coated 24-well plates (VWR). Prior infection, one well was used to determine the cell number by disrupting the suspension organoids into single cell suspension using TrypLE Express (Gibco) for 10-15min at 37°C and mechanical disruption if required. Organoids were either Mock infected in Advanced DMEM/F12 with 10 mM HEPES and 1xGlutaMax (all Thermo Fisher) (referred to as ADF++ medium) or challenged with respective IAV virus strain (MOI 1 or 10) for 1.5h at room temperature (RT). After infection, organoids were washed in ADF++ medium and resuspended in the respective culture medium without Gellan gum. Samples for flow cytometry, RT-qPCR, scRNA-seq and IAV replication analysis were taken at indicated time points.

### Flow cytometric analysis of infected LOs

To determine immunological response of infected organoids, Mock controls and infected samples were collected, dissociated into single cells and Fc blocked using 5μl Human TruStain FcX™ for 5min at RT. Samples were subsequently surface stained with antibodies anti-human EpCam (9C4),-HLA-DR (L243),-HLA-ABC (W6/32) from Biolegend for 30min at 4°C. Stained samples were fixed with FluoroFix Buffer (Biolegend) for 30min at 4°C and analyzed using a CytoFLEX flow cytometer (Beckman Coulter) and FlowJo-10 v10.8 software (Tree Star).

### Culture and infection of LOs in Gri3D plates

For single cell seeding of LOs into Gri3D 96 WP, hydrogel array IBIDI 400µm plates (SunBioscience, GRI3D-96IBI-S-96-400), liquid was removed via pipetting from the port and washed 2-3 times with respective LO medium and incubated for 30min at 37°C. Prior seeding, medium was removed first from the port and subsequently from the top of the well without touching the hydrogel. Single cell suspensions at densities of 60,000-150,000 cells/ml were pipetted as 50 µl drops directly onto the microwells. Single cells were let sediment for 30min at 37°C and gently over-layered with respective media supplemented with soluble 1-2% GFR Matrigel and 10 µM ROCKi via the port. Single LOs formed per microwell within 1-3 days and were expanded for 1-3 weeks until desired organoid size was reached before start of experiment. Matrigel content was sequentially diluted over time since it was not further supplemented after seeding. At the day of infection of LOs, cells from 3 wells were collected by disruption of the hydrogel using bored 200µl pipette tips. Collected organoids were retrieved out of the hydrogel and dissociated into single cells using TrypLE (Gibco) for 10min at 37°C. Single cell suspension was filtered through a 70µm cell strainer to get rid of excess hydrogel pieces and cells counted. For infection, medium was gently removed, and wells were gently washed 4x with 150µl either ADF++ medium only or supplemented with IAV (MOI 1 or 10). After 1.5h incubation at RT, all wells were washed again 4x with PBS^-/-^ before 4x addition of respective LO medium. For viral particle quantification, medium was sampled continuously from the same well. After sampling, medium was refilled and the dilution factor was included considered for calculation of total particle count. All bright field images were acquired using Leica DMi8 microscope.

### Plaque forming unit (PFU) assay

IAV infectious particles were quantified by plaque titration on MDCKII cells as previously described ^43,67^. Briefly, MDCKII monolayers at a confluency of 100% were incubated with virus-containing cell culture SNs at different dilutions for 45min at RT. After removal of remaining viral particles, infected cultures were gently overlaid with a 1:1 mixture of 2.5% Avicel medium and 2xMEM (Gibco) supplemented with 0.2% BSA (PAA), 0.001% Dextran, 0.05% NaHCO3 and 1 μg/ml TPCK-treated Trypsin (Sigma). After 48hpi, cells were washed twice with PBS^-/-^and plaques were fixed, visualized by staining with crystal violet and counted for viral particle quantification.

### Single-cell RNA Sequencing

At 24hpi, H3N2-infected apical-out iPSC-aLOs, along with apical-in and apical-out mock controls, were harvested and subsequently dissociated into single cell suspensions. Samples underwent Cell Multiplexing Oligo (CMO) labeling and pooling. Single cell capturing and downstream library construction were performed using the Chromium Single Cell 3′ V3.1 library preparation kit following the manufacturer’s instructions (10x Genomics). The sequencing libraries were multiplexed and loaded onto the flow cell of the Illumina® NovaSeq™ 6000 instrument following the manufacturer’s instructions. Sequencing was performed using a 2×150 paired-end configuration.

### Analysis of scRNA-seq data

Raw data was processed using Cellranger v6.0.0 and the GRCh38 reference augmented by the H3N2 IAV (DQ487333.1) genome. We then used CellBender to remove background RNA^68^. Data analysis was performed in R (version 4.2.2) with Seurat (v4.1.1). Cells with at least 250 detected genes, less than 20% mitochondrial or ribosomal content were combined from each library, library depth (total number of UMIs) and cell cycle scores were regressed out when scaling data. Datasets were then integrated using “Integrate Data”, followed by again regressing out library size, mitochondrial content, and cell cycle scores followed by doublet removal with DoubletFinder (version 2.0.3). A cluster of cells with high content of nuclear genes was removed, by means of a score for nuclear-localized genes using Seurat’s AddModuleScore and the top 50 genes enriched in single-nuc vs. single-cell libraries from Table S2 of Bakken et al ^69^. Data analysis was performed using CellXGene and plots were generated using R or CellxGene interactive Plugin. Gene Ontology analysis was performed using on-line tool https://go.princeton.edu/.

### Whole mount immunofluorescence staining

Apical-in and apical-out iPSC-LOs were fixed with 4% PFA for 1h at RT and washed two times with PBS. IF staining was performed in suspension in 1%BSA in PBS coated 1.5ml tubes using coated, bore tips. After fixation iPSC-LOs permeabilization and blocking were performed using 1%BSA, 5% donkey serum and 0.3%Triton X-100 in PBS for 1h at RT. Washes were performed using blocking buffer and soft centrifugation at 400rpm for 3min in less than 0.5ml. First antibody mouse anti-ZO-1 (33-9100, Thermo Fisher) was incubated overnight at 4°C. Secondary antibody donkey anti-mouse Alexa Fluor 647 (Thermo Fisher) was incubated for 2h at RT. Nuclei staining was done using 1µg/ml final concentration Dapi solution in PBS for 1h at RT. Images were acquired using Leica SP8 confocal microscope.

### Co-culture in Gri3D and live staining for high content imaging

For imaging, live organoids were stained in CK+DCI medium supplemented with 1:1000 SPY555-DNA (Spirochrome) one day prior to the co-culture. The staining solution was added through the port of the Gri3D plate, removed, and replenished 2–3 times to minimize dilution caused by incomplete volume exchange. TCPs were collected from G-Rex6M and rested overnight in complete medium in 24-well plates. On the day of co-culture, T cells were counted and stained with 10 µM eBioscience™ Cell Proliferation Dye eFluor™ 450 (Invitrogen) according to the manufacturer’s instructions. Prior addition of T cells, infection of organoids in Gri3D was performed as described above. After infection, organoids were washed 3x with 150µl PBS via the port and replenished with co-culture medium (1:1 mixture: T cell complete medium and cSFDM) containing 1:1000 Incucyte® Caspase-3/7 Reagent (red, Sartorius). The final volume was adjusted to 200µl per well. T cells were resuspended in co-culture medium supplemented with caspase-3/7 dye and 50,000 cells added in a 50µl drop directly on top of the respective wells of the Gri3D plate. After addition, T cells sedimented into the pockets of the respective microwells.

### Automated 3D high-content image acquisition and analysis of 3D T cell organoid co-culture

All images were acquired with the Opera Phenix High-Content Screening System (Revvity). Acquisition was performed in a two-step approach using the PreciScan feature. First, a lower magnification scan using an air objective (10x/ 0.3 NA) was used to pre-scan the complete well in XYZ-axis to identify positions of each organoid or region of interest (ROI) (Supplementary Table 3). ROIs were automatically selected and subsequently re-scanned at higher resolution using a water immersion objective (20x, 1.0 NA), one-peak autofocus and binning=2. Re-scan stacks were centered around at individual ROIs with 100-150 image planes taken 1µm apart. Acquisition of all fluorescent channels was separated. Images were analyzed using Harmony v5.1 software or Signals ImageArtist v1.3 (Revvity). Re-scanned stacks were analyzed in Harmony as a 3D volumes. Detailed analysis pipelines according to Supplementary Tables 4 (Low-resolution, pre-scan) and 5 (High-resolution, re-scan).

## Statistical analysis

Statistical analysis was performed using GraphPad Prism v10.3 (GraphPad Software). Data were first tested for normality using the Shapiro-Wilk and Kolmogorov-Smirnov tests. Depending on the experimental design, one-way or two-way ANOVA was then applied.

Statistical significance was set at p<0.05 and is indicated as follows: *p<0.05, **p<0.01, ***p<0.001, ****p<0.0001. Additional details specific to each dataset are provided in the corresponding figure legends.

## AI tools

During the preparation of the manuscript, ChatGPT (OpenAI) was used for optimization of the text and grammar correction. The model was used to support during the writing process, but all content was reviewed and finalized by the authors to ensure accuracy and originality. The authors of this manuscript retain full responsibility for the content of the manuscript.

## Data availability

Raw data will be deposited at [GEO Accession].

## Acknowledgements

We thank all voluntary blood donors for their donations. We thank Neşe Sevgili for performing RNA extractions and RT-qPCRs. We thank Toralf Kaiser and Jenny Kirsch from Flow Cytometry & Cell Sorting DRFZ for FACS sorting. We thank Anne Schwerk and Stephan Schlickeiser for critical discussion of statistical analysis. We thank Prof. D. Kotton for kindly providing human iPSC line BU3NG. We thank the CUSCO team for help with iPSC culture and full iPSC lines characterization. We thank Prof. Petra Reinke for financial and infrastructural support. Funding: the study was generously supported in parts by: Research Grant BCRT, BMBF TReAT 01EK2104A, Charité 3R. Cartoons in Figures were created with BioRender.com.

## Author information

### Contributions

U.D. designed and performed experiments, analyzed the data, composed the figures. V.FV. supervised, designed and performed experiments and analyzed data. U.D. and M.B. performed infection experiments and analyzed data. U.D. and N.W. produced TCPs and performed flow cytometric analysis. B.O. performed bioinformatic analysis of scRNA-seq data. A.K., V.FV., H.S. and U.D. designed and discussed image data analysis. T.F. supported experimental work with iPSC and organoid cultures. A.L., A.H., M.SH., V.FV., L.A. and H.S. supervised experiments. U.D., L.A. and H.S. developed and designed the project. U.D. and V.FV. wrote the manuscript with input from all co-authors. All co-authors approved the final version of the manuscript.

## Supplementary information

Supplementary figures 1-9 and Supplementary tables 1-5 are provided separately.

