## Supplementary Figures for "An autologous human iPSC-derived 3D organoid infection model for preclinical testing of antiviral T cells"

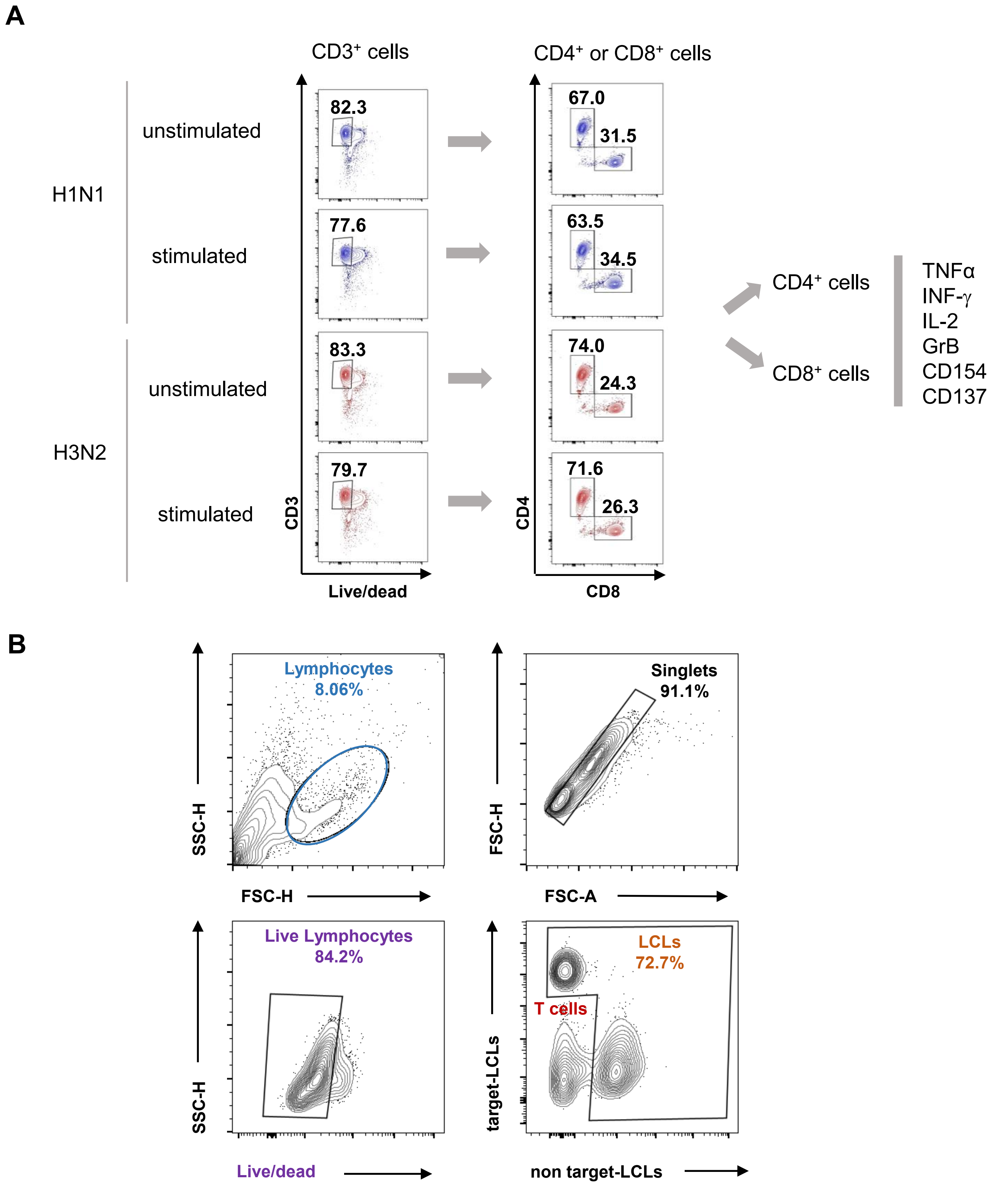


**Supplementary Figure 1. IAV-TCPs display effector functions *in vitro* and eliminate target-specific cells (flow cytometry gating strategies). A)** Representative flow cytometry plots showing gating strategy of unstimulated and peptide-stimulated H1N1- and H3N2-TCPs at d7 of expansion. CD4$^+$ and CD8$^+$ T cell populations were gated out of CD3$^+$ live cells. Production of effector cytokines TNF$\alpha$ , IFN-$\gamma$, IL-2, and molecule granzyme B (GrzB) and expression of activation markers CD154 and CD137 for each sub-population CD4$^+$ and CD8$^+$ are shown in Figure 2 A and B. **B)** Representative flow cytometry plots showing gating strategy of peptide-loaded target lymphoblastoid cell lines (LCLs) and non target-LCLs for killing assays. LCLs were gated out of live lymphocytes and T cell products were excluded from analysis to distinguish LCL populations. Killing assay results are shown in figure 2D.

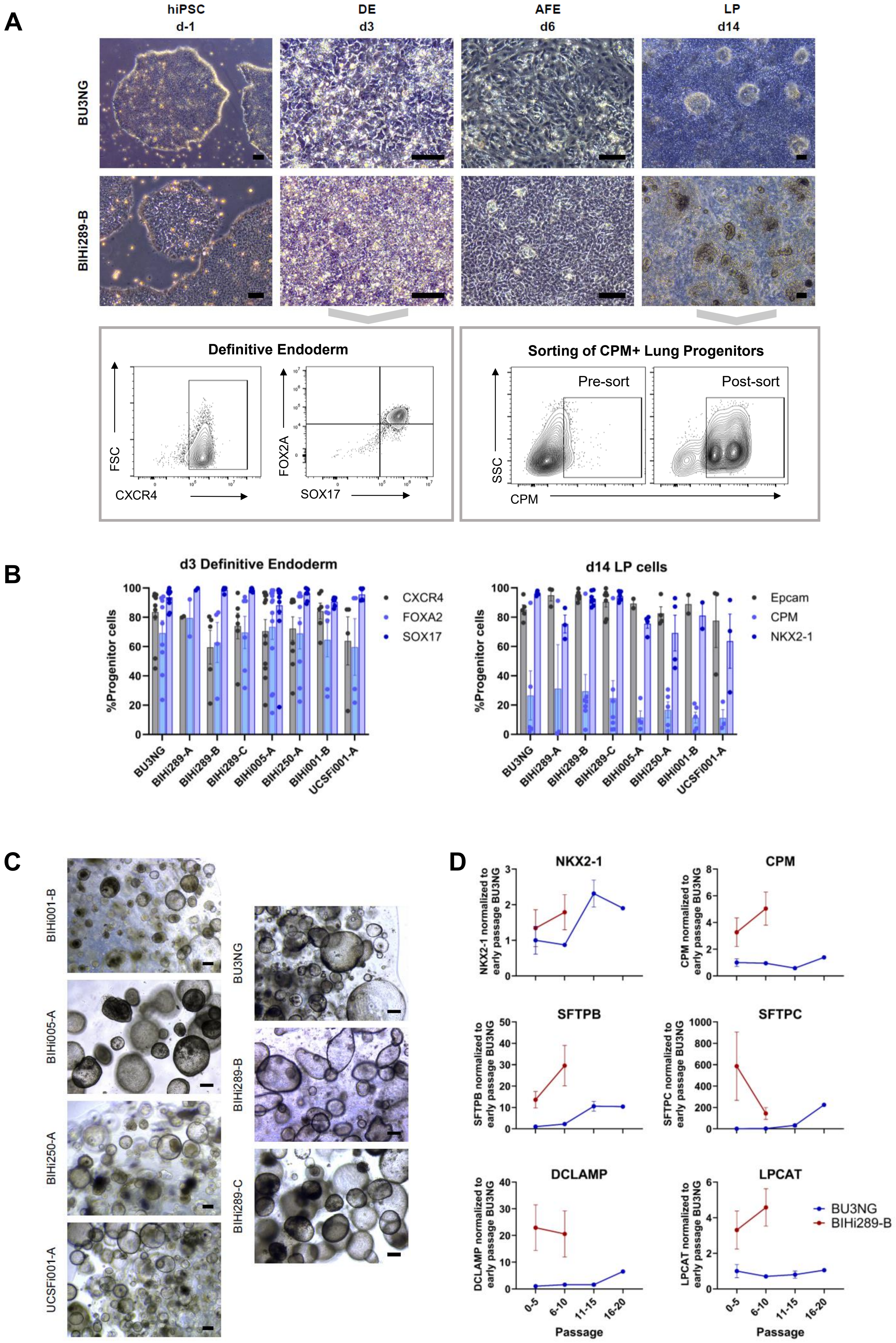

A
hiPSC d-1
DE d3
AFE d6
LP d14
BU3NG
BIHi289-B
Definitive Endoderm
FSC
CXCR4
FOX2A
SOX17
Sorting of CPM+ Lung Progenitors
Pre-sort
Post-sort
SSC
CPM
B
d3 Definitive Endoderm
%Progenitor cells
CXCR4
FOXA2
SOX17
d14 LP cells
Epcam
CPM
NKX2-1
BU3NG
BIHi289-A
BIHi289-B
BIHi289-C
BIHi005-A
BIHi250-A
BIHi001-B
UCSFi001-A
C
BIHi001-B
BIHi005-A
BIHi250-A
UCSFi001-A
BU3NG
BIHi289-B
BIHi289-C
D
NKX2-1
NKX2-1 normalized to early passage BU3NG
CPM
CPM normalized to early passage BU3NG
SFTPB
SFTPB normalized to early passage BU3NG
SFTPC
SFTPC normalized to early passage BU3NG
DCLAMP
DCLAMP normalized to early passage BU3NG
LPCAT
LPCAT normalized to early passage BU3NG
Passage
0-5
6-10
11-15
16-20
BU3NG
BIHi289-B

**Supplementary Figure 2. iPSC-aLOs differentiation is robust across hiPSC lines. A)** Timeline with representative images of iPSC differentiation to LP with representative images at each stage (upper panel). Representative flow plots showing gating strategy for quality assessment of DE differentiation at d3 and sorting efficiency of carboxypeptidase M (CPM) for LPs. **B)** DE (left) and LP (right) differentiation efficiencies quantified by the frequency of $CXCR4^{+}$, $FOXA2^{+}$, $SOX17^{+}$ cells and Epcam+, $CPM^{+}$, $NKX2\text{-}1^{+}$ cells, respectively. Results are expressed as mean ±SEM (n=2-13). **C)** Representative images of iPSC-aLOs generated from different iPSC-lines. Scale bars: 200μm **D)** Real time quantitative PCR results showing lung markers expressed in iPSC-aLOs derived from different iPSC lines across passages. Gene expression is normalized to earlier passage of line BU3NG (used as reference in this study). Results are expressed as mean ±SEM (n=2-9). ***DE:*** *definitive endoderm,* ***AFE:*** *anterior-foregut endoderm,* ***LP:*** *lung progenitors.*

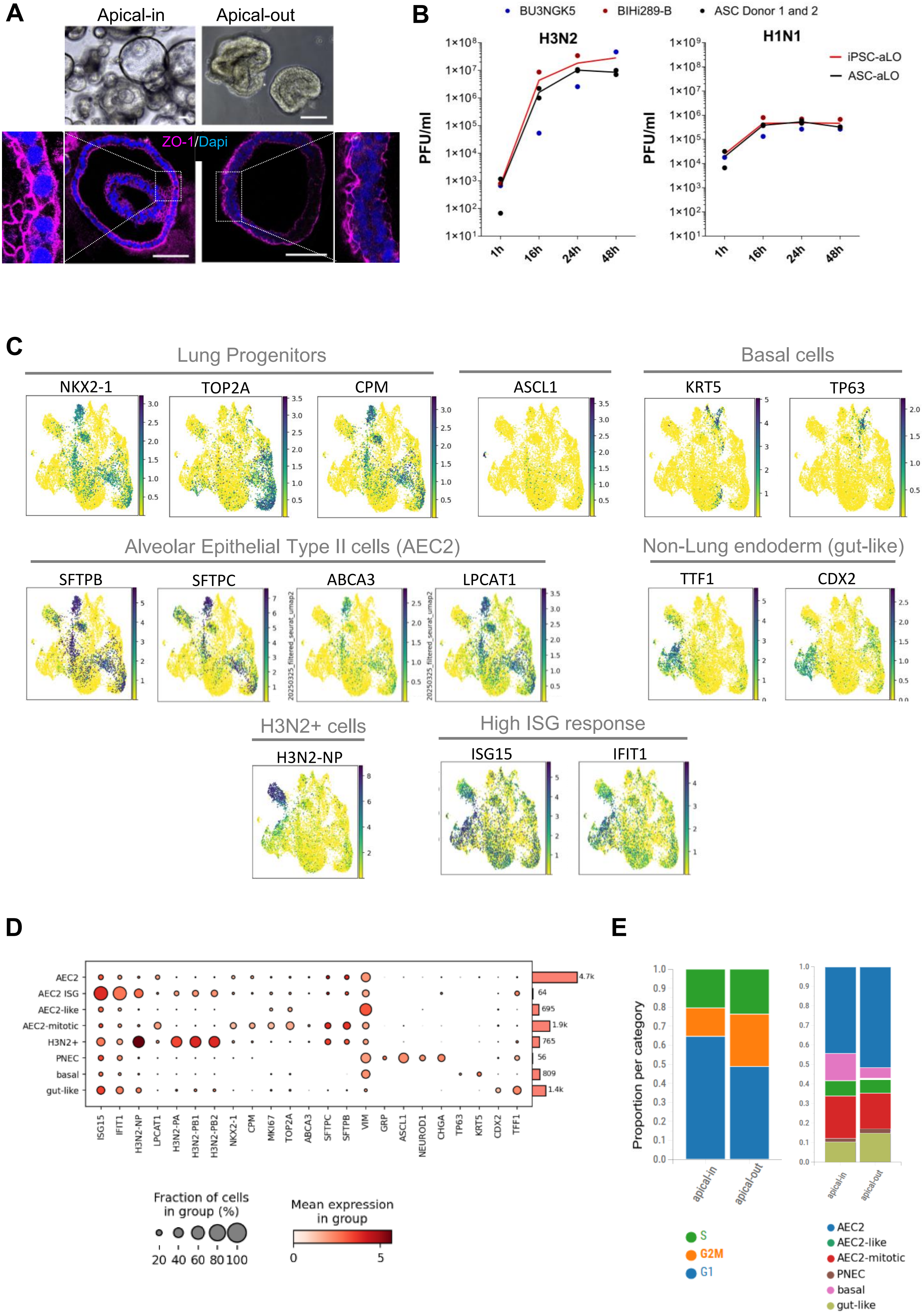
A
Apical-in
Apical-out
ZO-1/Dapi
B
BU3NGK5
BIHi289-B
ASC Donor 1 and 2
H3N2
H1N1
PFU/ml
iPSC-aLO
ASC-aLO
1h
16h
24h
48h
C
Lung Progenitors
NKX2-1
TOP2A
CPM
ASCL1
Basal cells
KRT5
TP63
Alveolar Epithelial Type II cells (AEC2)
SFTPB
SFTPC
ABCA3
LPCAT1
Non-Lung endoderm (gut-like)
TTF1
CDX2
H3N2+ cells
H3N2-NP
High ISG response
ISG15
IFIT1
D
AEC2
AEC2 ISG
AEC2-like
AEC2-mitotic
H3N2+
PNEC
basal
gut-like
4.7k
64
695
1.9k
765
56
809
1.4k
ISG15
IFIT1
H3N2-NP
LPCAT1
H3N2-PA
H3N2-PB1
H3N2-PB2
NKX2-1
CPM
MKI67
TOP2A
ABCA3
SFTPC
SFTPB
VIM
GRP
ASCL1
NEUROD1
CHGA
TP63
KRT5
CDX2
TFF1
Fraction of cells
in group (%)
20 40 60 80 100
Mean expression
in group
0
5
E
Proportion per category
apical-in
apical-out
S
G2M
G1
AEC2
AEC2-like
AEC2-mitotic
PNEC
basal
gut-like

**Supplementary Figure 3. iPSC-aLOs phenotypic characteristics**. **A)** Representative bright field and immunofluorescence images of iPSC-aLOs in apical-out and apical-in conformation. Scale bars: 100µm. **B)** Quantification of plaque forming units (PFU), replication curves are shown for iPSC-aLOs (n=2 independent hiPSC lines) and ASC-aLOs (n=2) upon H3N2 and H1N1 infection up to 48hpi. **C)** UMAPs showing cluster marker gene expression according to annotated populations. **D)** Dot plot showing marker genes expression per cluster of single-cell RNA sequencing of H3N2-infected and uninfected iPSC-aLOs at 24hpi (n=2, hiPSC lines as biological replicates). **E)** Cell cycle phase (left) and cell type cluster (right) distribution for apical-in and apical-out iPSC-aLOs conformation in control samples (uninfected). ***iPSC-aLOs:*** *hiPSC-derived alveolar-like lung organoids.*

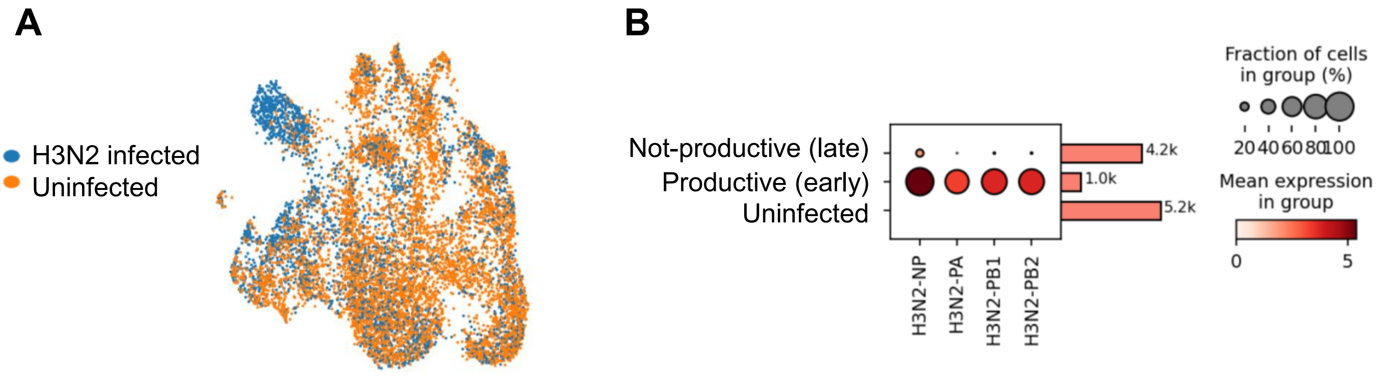


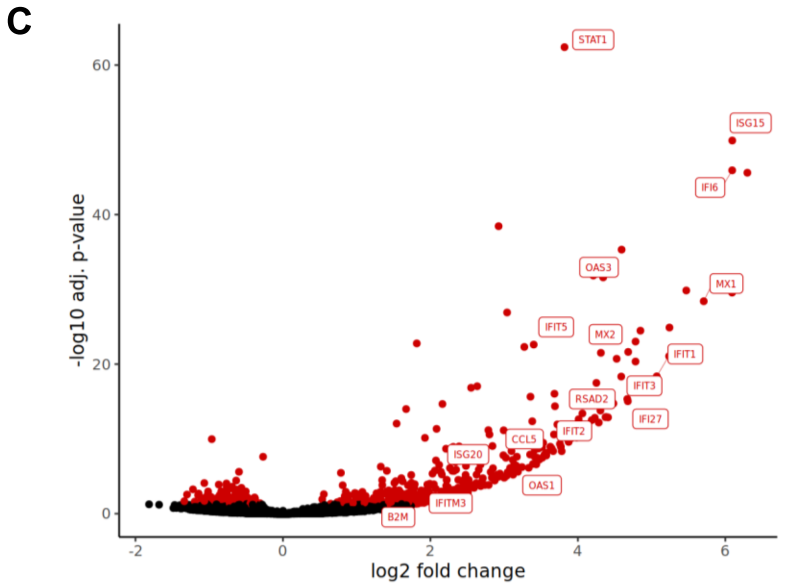


| GO term | p-Value |
|---|---|
| Defense response to virus | 3.88E-63 |
| Innate immune response | 5.92E-51 |
| Immune response | 4.58E-46 |
| Regulation of viral process | 9.98E-34 |
| Response to cytokine | 1.98E-27 |
| Viral genome replication | 8.99E-33 |
| Interferon-mediated signaling pathway | 6.54E-26 |
| Response to type I interferon | 1.86E-22 |
| Viral process | 6.89E-22 |
| Viral life cycle | 5.04E-19 |
| Response to type II interferon | 6.67E-15 |
| Antiviral innate immune response | 4.83E-15 |
| T cell mediated immunity | 1.40E-10 |
| Cell killing | 1.58E-09 |
| Antigen processing and presentation of peptide antigen via MHC class I | 1.18E-05 |

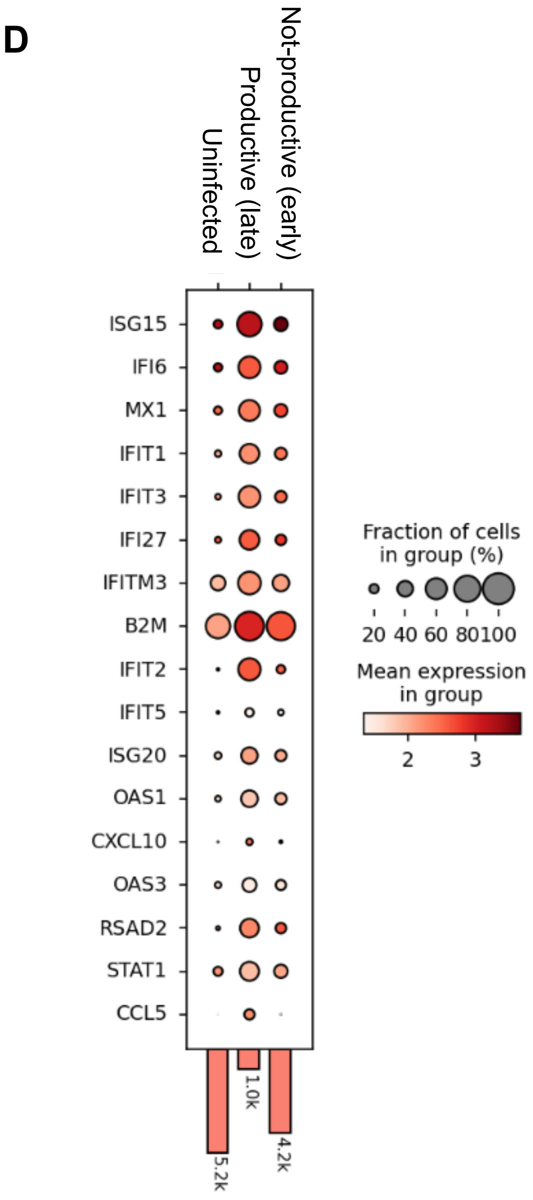


**Supplementary Figure 4. iPSC-aLOs show interferon stimulated gene response upon H3N2 infection. A)** UMAP showing distribution of infected and uninfected cells. **B)** Dot plot showing expression of viral genes and fraction of cells per group for infected, likely productive infected and uninfected cells by single-cell RNA sequencing (scRNA-seq). **C)** Vulcano plot showing differential gene expression of infected versus uninfected samples from scRNA-seq with interferon stimulated genes (ISGs) highlighted (upper panel). Gene ontology (GO) terms given for up-regulated genes (lower panel). **D)** Dot plot indicating expression levels of ISG among infected, likely productive and uninfected single cell clusters. ***iPSC-aLOs:*** *hiPSC-derived alveolar-like lung organoids;* ***ASC-aLOs:*** *adult stem cell-derived alveolar-like lung organoids.*

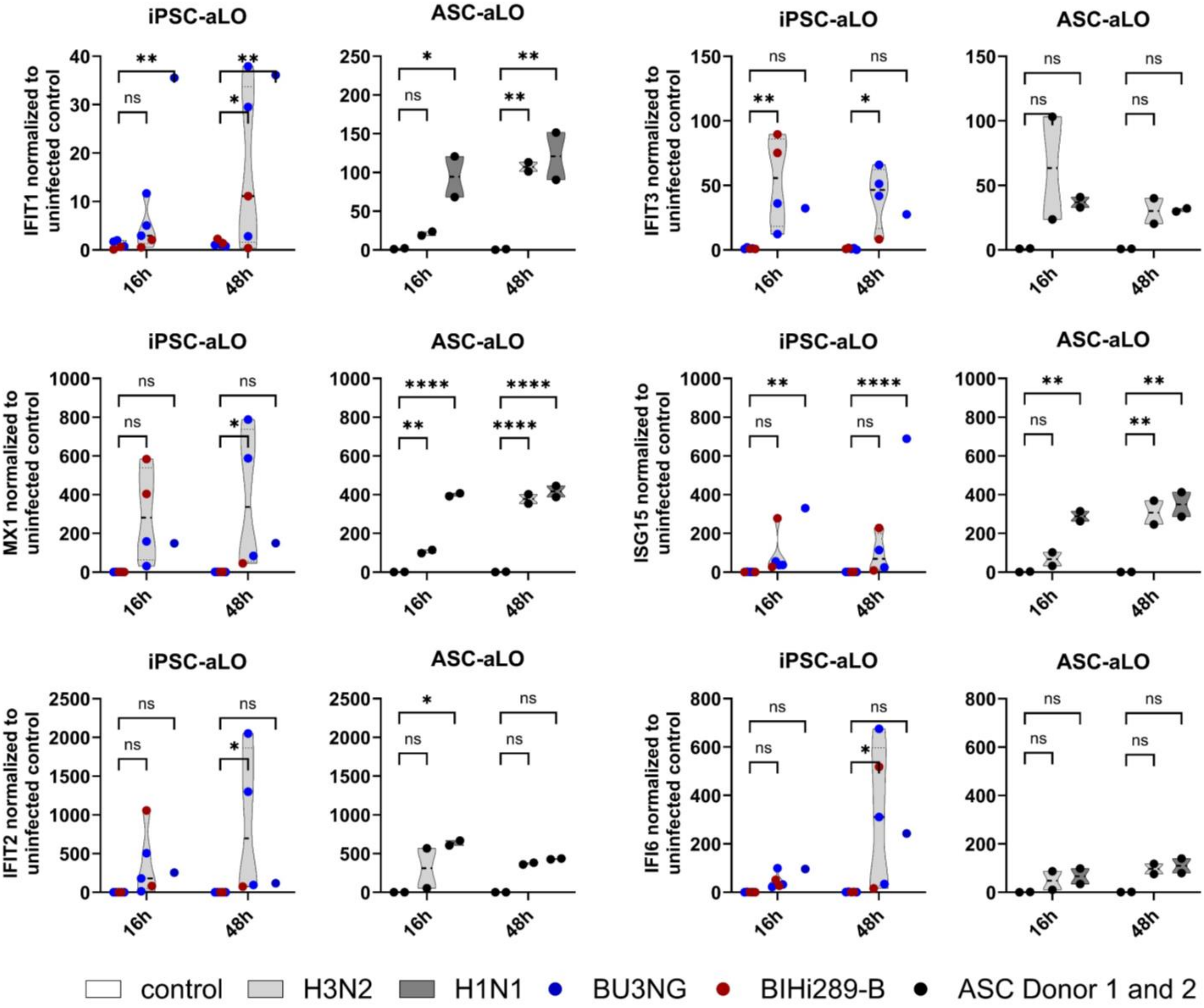


**Supplementary Figure 5. IAV infection triggers similar responses in iPSC-aLOs and ASC-aLOs.** Real time quantitative PCR results showing expression of interferon stimulated genes in H3N2- and H1N1-infected iPSC-aLOs and ASC-aLOs at 16 and 48 hours post infection. Gene expression is normalized to uninfected control per experiment. Each dot represents individual donors. Experiments with the same donor were run as independent replicates. Two-way ANOVA Kruskal-Wallis Test p<0.05 (*), p<0.01 (**), p<0.001 (***), p<0.0001 (****). ***iPSC-aLOs:*** *hiPSC derived alveolar-like lung organoids;* ***ASC-aLOs:*** *adult stem cell derived alveolar-like lung organoids.*

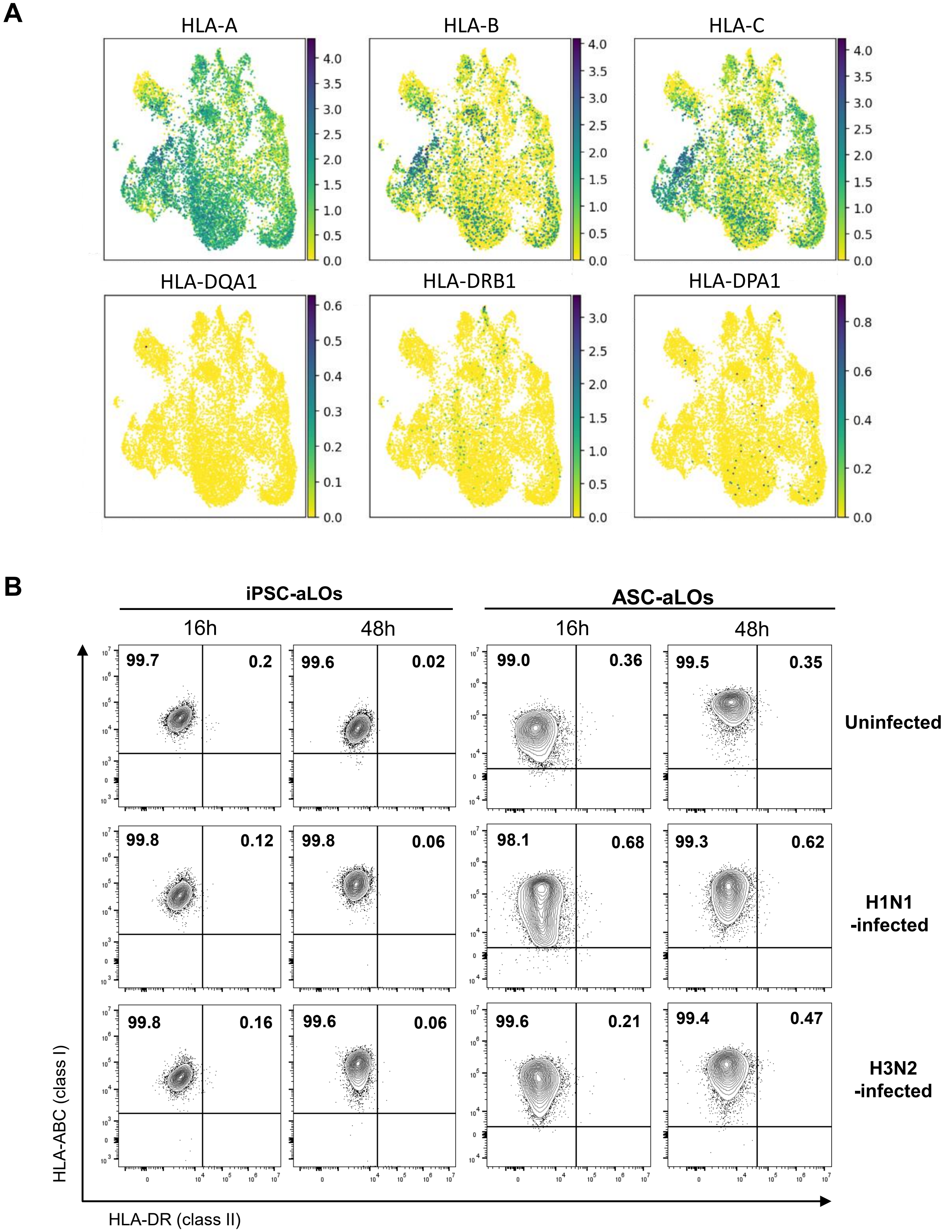


**Supplementary Figure 6: iPSC-aLOs express HLA class I. A)** UMAPs derived from single-cell RNA sequencing analysis showing HLA class I and class II subtype expression in iPSC-aLOs 24hs after infection (all cells are shown). **B)** Representative flow cytometric plots showing HLA class I (ABC) and HLA class II (DR-subtype) expression on iPSC-aLOs and ASC-aLOs at indicated timepoint post infection with H3N2 or H1N1.

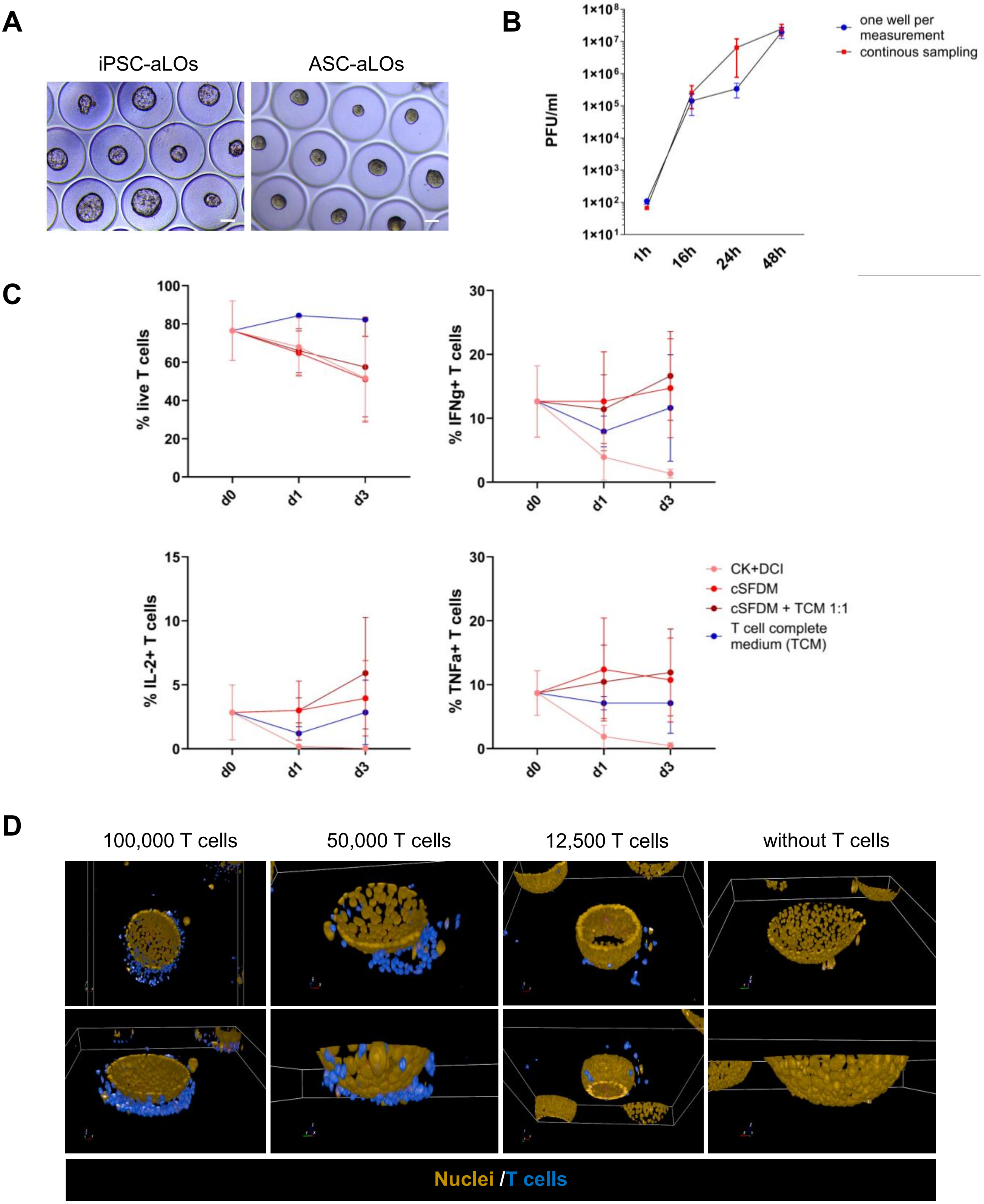


**Supplementary Figure 7. Lung organoids and T cells co-culture establishment. A)** Representative bright field images showing size of iPSC-aLOs and ASC-aLOs cultured in Gri3D prior infection. Scale bar: 100 µm. **B)** Virus particle quantification using plaque forming unit (PFU) assay of H3N2-infected ASC-aLOs in Gri3D. Results are shown as mean ±SEM (n=3). Sampling comparison: one well per timepoint versus continuous sampling from same well. **C)** Flow cytometric assessment showing T cell viability and frequency of effector cytokine production over time for different co-culture formulations: iPSC-aLOs complete medium (CK+DCI), iPSC-aLOs basal medium (cSFDM), cSFDM plus T cell complete medium (TCM) 1:1 and TCM alone. Results are shown as mean ±SEM (n=2). **D)** Representative 3D images showing titration of T cells added to iPSC-aLOs for subsequent segmentation. ***iPSC-aLOs:*** *hiPSC derived alveolar-like lung organoids;* ***ASC-aLOs:*** *adult stem cell derived alveolar-like lung organoids.*

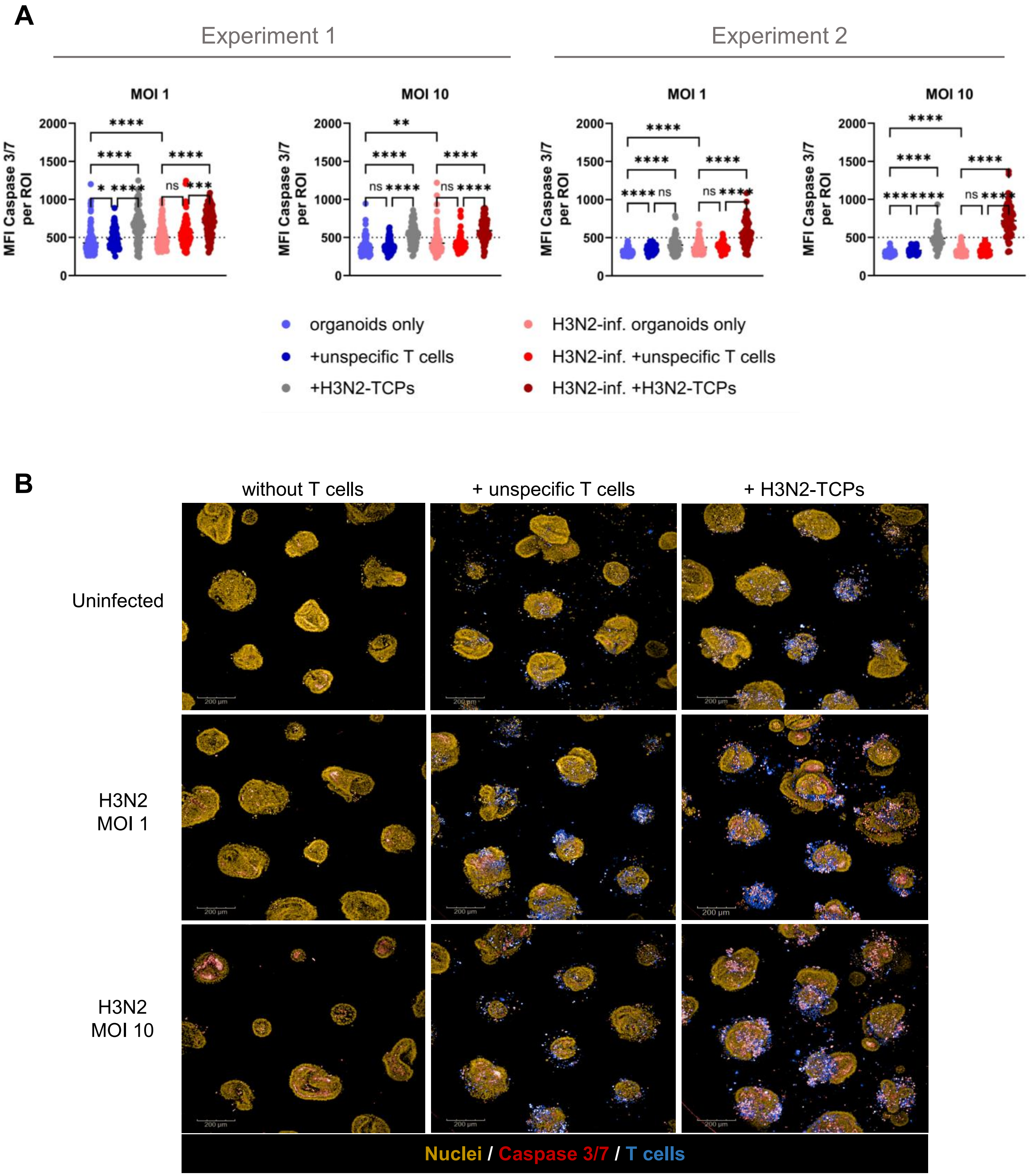


**Supplementary Figure 8: Robustness of imaging assay at low resolution. A)** Plots showing mean fluorescence intensity (MFI) for Caspase 3/7$^{+}$ quantification at endpoint analysis of H3N2-infected iPSC-aLOs in co-culture with autologous T cells (48 hours post infection). Two for 2 biological replicates (n= 1 donor, n= 2 independent infection and co-culture experiments) using 2 different MOI (1 and 10) are shown separately. Dots represent each region of interest (ROI) composed by one iPSC-aLO + T cells. Two-way ANOVA Kruskal-Wallis Test p<0.05 (*), p<0.01 (**), p<0.001 (***), p<0.0001 (****). **B)** Representative panel showing fluorescence images acquired at low magnification (pre-scan) for H3N2-infected and uninfected iPSC-aLOs in co-culture with or without T cells (unspecific and H3N2-TCP). Scale bars: 200µm. ***iPSC-aLO:** hiPSC derived alveolar-like lung organoid*, ***TCP**: T cell product.*

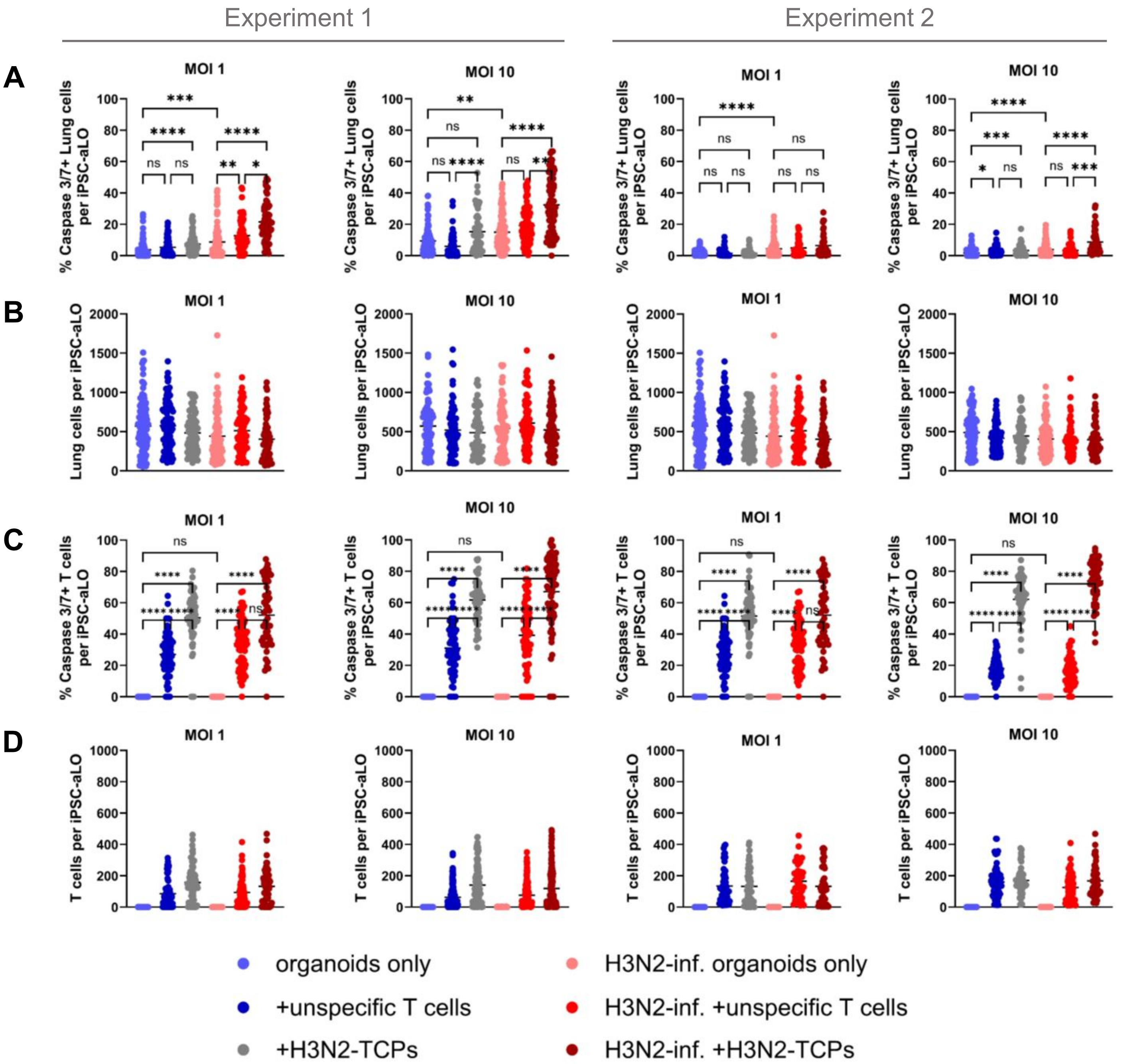


**Supplementary Figure 9. Robustness of imaging assay at high resolution.** Endpoint analysis of H3N2-infected iPSC-aLOs in co-culture with autologous T cells at 48 hours post infection is shown separately for 2 biological replicates (n= 1 donor, n= 2 independent infection and co-culture experiments) using 2 different MOIs (1 and 10). Dots represent each iPSC-aLO. **A)** Plot showing percentage of Caspase 3/7$^{+}$ lung cells per organoid (iPSC-aLO). **B)** Total number of quantified lung cells per organoid (iPSC-aLO). **C)** Plot showing percentage of Caspase 3/7$^{+}$ T cells per organoid (iPSC-aLO). **D)** Total number of quantified T cells per organoid (iPSC-aLO). Two-way ANOVA Kruskal-Wallis Test p<0.05 (*), p<0.01 (**), p<0.001 (***), p<0.0001 (****).

***iPSC-aLO:*** *hiPSC derived alveolar-like lung organoid.*
