## Supplementary Table 3 for "An autologous human iPSC-derived 3D organoid infection model for preclinical testing of antiviral T cells"

### Analysis Sequence "PreScan_Analysis XYZ Gri3D"

| Input Image | Input |
|---|---|
| | **Channel group :** 1<br>**Sequences :** ALL<br>**Flatfield Correction :** None<br>Brightfield Correction<br>**Stack Processing :** Maximum Projection<br>Create Global Image<br>**Min. Global Binning :** Dynamic |

| Filter Image | Input | Method | Output |
|---|---|---|---|
| | **Channel :** SPY555-DNA (global) | **Method :** Sliding Parabola<br>Curvature : 8 | Output Image : Sliding Parabola (global) |

| Find Image Region | Input | Method | Output |
|---|---|---|---|
| | **Channel :** Sliding Parabola (global)<br>**ROI :** Imaged Area (global)<br>**ROI Region :** Imaged Area | **Method :** Whole Image Region | Output Population : Whole Image (global)<br>Output Region : Whole Image Region |

| Select Region | Input | Method | Output |
|---|---|---|---|
| | **Population :** Whole Image (global)<br>**Region :** Whole Image Region | **Method :** Resize Region [%]<br>Outer Border : 5 %<br>Outer Population : None<br>Outer Region :<br>Inner Border : 100 % | Output Region : Resized Region |

| Find Image Region (2) | Input | Method | Output |
|---|---|---|---|
| | **Channel :** Sliding Parabola (global)<br>**ROI :** Whole Image (global)<br>**ROI Region :** Resized Region | **Method :** Absolute Threshold<br>Lowest Intensity : ≥ 300<br>Highest Intensity : ≤ INF<br>Split into Objects<br>Area : > 300 µm²<br>Fill Holes | Output Population : ROI Candidate<br>Output Region : ROI |

| Modify Population | Input | Method | Output |
|---|---|---|---|
| | **Population :** ROI Candidate<br>**Region :** Organoid | **Method :** Cluster by Distance<br>Distance : 2 px<br>Area : > 0 px² | Output Population : ROI Candidate Clustered<br>Output Region : ROI Candidate Clustered Region |

| Calculate Intensity Properties | Input | Method | Output |
|---|---|---|---|
| | **Channel :** SPY555-DNA (global)<br>**Population :** ROI Candidate Clustered<br>**Region :** ROI Candidate Clustered Region | **Method :** Standard Mean | Property Prefix : ROI Candidate Clustered Intensity SPY555 |

| Calculate Morphology Properties | Input | Method | Output |
|---|---|---|---|
| | **Population :** ROI Candidate Clustered<br>**Region :** ROI Candidate Clustered Region | **Method :** Standard (deprecated)<br>Area<br>Roundness<br>Width | Property Prefix : ROI Clustered Morphology |

| Select Population | Input | Method | Output |
|---|---|---|---|
| | **Population :** ROI Candidate Clustered | **Method :** Filter by Property<br>ROI Clustered Morphology Roundness : > 0.2<br>ROI Clustered Morphology Area [µm²] : > 2000<br>ROI Clustered Morphology Area [µm²] : < 100000<br>Boolean Operations : F1 and F2 and F3 | Output Population : ROI Selected |

| Determine Well Layout ⚠ | Input | Method | Output |
|---|---|---|---|
| | **Population :** ROI Selected<br>**Region :** ROI Candidate Clustered Region | **Method XY :** Individual Objects<br>Rescan Magnification : 20x<br>Rescan Camera ROI : 2160x2160<br>Max No of Fields :<br>Object Margin : 20 µm<br>Field Overlap : 2 %<br><br>**Method Z :** By Plane Map<br>Plane Map : Plane Map SPY555-DNA (global) | Output Population : Well Layout (global) |

| Define Results ⚠ | Results |
|---|---|
| | **Method :** Standard Output<br>Clustered Organoid Candidate - Number of Objects : Object Count<br>Output Name : Clustered Organoid Candidate - Number of Objects<br><br>**Method :** List of Outputs |

**Object Results**

Population : Whole Image (global) : None
Population : Clustered Organoid Candidate : None
Population : Organoid Candidate : None
Population : Clustered Organoid Candidate Selected : None

Acapella version: 5.4.1.131587. Timestamp: 2025-05-07 12:47:46 +0200.
