## Supplementary Table 4 for "An autologous human iPSC-derived 3D organoid infection model for preclinical testing of antiviral T cells"

### Analysis Sequence "Low resolution downstream analysis"

| Input Image | Input |
|---|---|
| | **Channel group :** 1<br>**Sequences :** ALL<br>**Flatfield Correction :** Basic<br>Brightfield Correction<br>**Stack Processing :** Maximum Projection<br>Create Global Image<br>**Min. Global Binning :** Dynamic |

| Find Image Region | Input | Method | Output |
|---|---|---|---|
| | **Channel :** SPY555-DNA (global)<br>**ROI :** Imaged Area (global)<br>**ROI Region :** Imaged Area | **Method :** Common Threshold<br>Threshold : 0.5<br>Split into Objects<br>Area : > 3000 µm²<br>Fill Holes | Output Population : ROI 1<br>Output Region : Image Region |

| Select Region | Input | Method | Output |
|---|---|---|---|
| | **Population :** ROI 1<br>**Region :** Image Region | **Method :** Resize Region [%]<br>Outer Border : 42 %<br>Outer Population : Imaged Area (global)<br>Outer Region : Imaged Area<br>Inner Border : 100 % | Output Region : ROI 1 Resized |

| Calculate Texture Properties | Input | Method | Output |
|---|---|---|---|
| | **Channel :** SPY555-DNA (global)<br>**Population :** ROI 1<br>**Region :** ROI 1 Resized | **Method :** SER Features<br>Scale : 0 px<br>Normalization by : Kernel<br>SER Spot<br>SER Hole<br>SER Edge<br>SER Ridge<br>SER Valley<br>SER Saddle<br>SER Bright<br>SER Dark | Property Prefix : ROI 1 Resized SPY555 |

| Calculate Texture Properties (2) | Input | Method | Output |
|---|---|---|---|
| | **Channel :** SPY555-DNA (global)<br>**Population :** ROI 1<br>**Region :** ROI 1 Resized | **Method :** Haralick Features<br>Distance : 1 px<br>Haralick Contrast | Property Prefix : ROI 1 Resized SPY555 |

| | | |
|---|---|---|
| | | Haralick Correlation<br>Haralick Sum Variance<br>Haralick Homogeneity |

| **Calculate Texture Properties (3)** | **Input** | **Method** | **Output** |
|---|---|---|---|
| | **Channel :** SPY555-DNA (global)<br>**Population :** ROI 1<br>**Region :** ROI 1 Resized | **Method :** Gabor Features<br>Scale : 2 px<br>Wavelength : 2<br>Number of Angles : 8<br>Normalization by : Kernel<br>Gabor Min<br>Gabor Max | Property Prefix : ROI 1 Resized SPY555-DNA (global) |

| **Calculate Texture Properties (4)** | **Input** | **Method** | **Output** |
|---|---|---|---|
| | **Channel :** ef450 (global)<br>**Population :** ROI 1<br>**Region :** ROI 1 Resized | **Method :** SER Features<br>Scale : 0 px<br>Normalization by : Kernel<br>SER Spot<br>SER Hole<br>SER Edge<br>SER Ridge<br>SER Valley<br>SER Saddle<br>SER Bright<br>SER Dark | Property Prefix : ROI 1 Resized ef450 (global) |

| **Calculate Texture Properties (5)** | **Input** | **Method** | **Output** |
|---|---|---|---|
| | **Channel :** ef450 (global)<br>**Population :** ROI 1<br>**Region :** ROI 1 Resized | **Method :** Haralick Features<br>Distance : 1 px<br>Haralick Contrast<br>Haralick Correlation<br>Haralick Sum Variance<br>Haralick Homogeneity | Property Prefix : ROI 1 Resized ef450 (global) |

| **Calculate Texture Properties (6)** | **Input** | **Method** | **Output** |
|---|---|---|---|
| | **Channel :** ef450 (global)<br>**Population :** ROI 1<br>**Region :** ROI 1 Resized | **Method :** Gabor Features<br>Scale : 2 px<br>Wavelength : 2<br>Number of Angles : 8<br>Normalization by : Kernel<br>Gabor Min<br>Gabor Max | Property Prefix : ROI 1 Resized ef450 (global) |

| **Calculate Intensity Properties** | **Input** | **Method** | **Output** |
|---|---|---|---|

| | | | |
|---|---|---|---|
| | **Channel :** SPY555-DNA (global)<br>**Population :** ROI 1<br>**Region :** ROI 1 Resized | **Method :** Standard Mean | Property Prefix : Intensity ROI 1 Resized SPY555-DNA (global) |
| **Calculate Intensity Properties (2)** | **Input** | **Method** | **Output** |
| | **Channel :** ef450 (global)<br>**Population :** ROI 1<br>**Region :** ROI 1 Resized | **Method :** Standard Mean | Property Prefix : Intensity ROI 1 Resized ef450 (global) |
| **Select Population** | **Input** | **Method** | **Output** |
| | **Population :** ROI 1 | **Method :** Linear Classifier<br>Number of Classes : 2<br>Organoid Region Resized SPY555-DNA (global) SER Spot 0 px<br>Organoid Region Resized SPY555-DNA (global) SER Hole 0 px<br>Organoid Region Resized SPY555-DNA (global) SER Edge 0 px<br>Organoid Region Resized SPY555-DNA (global) SER Ridge 0 px<br>Organoid Region Resized SPY555-DNA (global) SER Valley 0 px<br>Organoid Region Resized SPY555-DNA (global) SER Saddle 0 px<br>Organoid Region Resized SPY555-DNA (global) SER Bright 0 px<br>Organoid Region Resized SPY555-DNA (global) SER Dark 0 px<br>Organoid Region Resized SPY555-DNA (global) Haralick Correlation 1 px<br>Organoid Region Resized SPY555-DNA (global) Haralick Contrast 1 px<br>Organoid Region Resized SPY555-DNA (global) Haralick Sum Variance 1 px<br>Organoid Region Resized SPY555-DNA (global) Haralick Homogeneity 1 px<br>Organoid Region Resized SPY555-DNA (global) Gabor Min 2 px w2<br>Organoid Region Resized SPY555-DNA (global) Gabor Max 2 px w2 | Output Population A : iPSC-aLO + T cells<br>Output Population B : T cells only |

| | | |
|---|---|---|
| | | Image Region ef450 (global) SER Spot 0 px<br>Image Region ef450 (global) SER Hole 0 px<br>Image Region ef450 (global) SER Edge 0 px<br>Image Region ef450 (global) SER Ridge 0 px<br>Image Region ef450 (global) SER Valley 0 px<br>Image Region ef450 (global) SER Saddle 0 px<br>Image Region ef450 (global) SER Bright 0 px<br>Image Region ef450 (global) SER Dark 0 px<br>Organoid Region Resized ef450 (global) Haralick Correlation 1 px<br>Organoid Region Resized ef450 (global) Haralick Contrast 1 px<br>Organoid Region Resized ef450 (global) Haralick Sum Variance 1 px<br>Organoid Region Resized ef450 (global) Haralick Homogeneity 1 px<br>Organoid Region Resized ef450 (global) Gabor Min 2 px w2<br>Organoid Region Resized ef450 (global) Gabor Max 2 px w2<br>Intensity Organoid Region Resized SPY555-DNA (global) Mean<br>Intensity Organoid Region Resized ef450 (global) Mean |

| **Calculate Intensity Properties (4)** | **Input** | **Method** | **Output** |
|---|---|---|---|
| | **Channel :** Alexa 647 (global)<br>**Population :** iPSC-aLO + T cells<br>**Region :** ROI 1 Resized | **Method :** Standard Mean | Property Prefix : Intensity Casp iPSC-aLO + T cells |

| **Calculate Properties** | **Input** | **Method** | **Output** |
|---|---|---|---|
| | **Population :** ROI 1 | **Method :** By Related Population<br>Related Population : iPSC-aLO + T cells<br>Intensity Casp iPSC-aLO + T cells Mean : Mean | Property Suffix : per Object |

| **Define Results** | **Results** |
|---|---|

**Method :** List of Outputs
**Population : iPSC-aLO + T cells**
Intensity Casp iPSC-aLO + T cells Mean : Mean

**Population : ROI 1**
Number of Objects

**Object Results**
Population : T cells only : None
Population : iPSC-aLO + T cells : Use Selected Well Results
Population : ROI 1 : Use Selected Well Results

Acapella version: 5.4.1.131587. Timestamp: 2025-05-07 11:54:57 +0200.
