## Supplementary Table 5 for "An autologous human iPSC-derived 3D organoid infection model for preclinical testing of antiviral T cells"

### Analysis Sequence "High resolution downstream analysis"

| Input Image | Input |
|---|---|
| | **Channel group :** 1<br>**Sequences :** ALL<br>**Flatfield Correction :** Basic Brightfield Correction<br>**Stack Processing :** 3D Analysis |

| Find Image Region | Input | Method | Output |
|---|---|---|---|
| | **Channel :** SPY555-DNA<br>**ROI :** None | **Method :** Local Threshold<br>Threshold : 0.5<br>Region Scale : 10 µm<br>Join touching Fragments<br>Volume : > 100000 µm³ | Output Population : Image Region<br>Output Region : Image Region |

| Select Population | Input | Method | Output |
|---|---|---|---|
| | **Population :** Image Region | **Method :** Common Filters<br>Remove Objects : Touching Side Faces<br>Region : Image Region | Output Population : Image Region 2 |

| Modify Population | Input | Method | Output |
|---|---|---|---|
| | **Population :** Image Region 2<br>**Region :** Image Region | **Method :** Cluster by Distance<br>Distance : 10 µm<br>Volume : > 0 µm³ | Output Population : Organoid<br>Output Region : Organoid |

| Filter Image | Input | Method | Output |
|---|---|---|---|
| | **Channel :** SPY555-DNA | **Method :** Smoothing<br>Filter : Gaussian<br>Width : 5 µm | Output Image : Gaussian Smoothed |

| Calculate Image | Input | Method | Output |
|---|---|---|---|
| | | **Method :** By Formula<br>Formula : A-B<br>Channel A : SPY555-DNA<br>Channel B : Gaussian Smoothed<br>Negative Values : Set to Zero<br>Undefined Values : Set to Zero | Output Image : Calculated Image |

| Filter Image (2) | Input | Method | Output |
|---|---|---|---|

| | | | |
|---|---|---|---|
| | **Channel :** Calculated Image | **Method :** Smoothing<br>Filter : Gaussian<br>Width : 1 px | Output Image : Gaussian Smoothed (2) |

| Calculate Image (2) | Input | Method | Output |
|---|---|---|---|
| | | **Method :** By Formula<br>Formula : A*2+B<br>Channel A : Calculated Image<br>Channel B : Gaussian Smoothed (2)<br>Negative Values : Set to Zero<br>Undefined Values : Set to Zero | Output Image : Calculated Image (2) |

| Find Nuclei | Input | Method | Output |
|---|---|---|---|
| | **Channel :** Calculated Image<br>**ROI :** Organoid<br>**ROI Region :** Organoid | **Method :** C<br>Common Threshold : 0<br>Volume : > 150 µm³<br>Splitting Coefficient : 2.8<br>Individual Threshold : 0.05<br>Contrast : > -1<br>Accuracy / Speed : Standard / Standard | Output Population : Nuclei |

| Calculate Intensity Properties (3) | Input | Method | Output |
|---|---|---|---|
| | **Channel :** SPY555-DNA<br>**Population :** Nuclei<br>**Region :** Nucleus | **Method :** Standard Mean | Property Prefix : Intensity SPY555 Nuclei |

| Calculate Intensity Properties | Input | Method | Output |
|---|---|---|---|
| | **Channel :** ef450<br>**Population :** Nuclei<br>**Region :** Nucleus | **Method :** Standard Mean | Property Prefix : Intensity ef450 Nuclei |

| Calculate Intensity Properties (2) | Input | Method | Output |
|---|---|---|---|
| | **Channel :** Alexa 647<br>**Population :** Nuclei<br>**Region :** Nucleus | **Method :** Standard Mean | Property Prefix : Intensity AF647 Nuclei |

| Select Population (2) | Input | Method | Output |
|---|---|---|---|

| | | | |
|---|---|---|---|
| | **Population :** Nuclei | **Method :** Filter by Property<br>Intensity ef450 Nuclei Mean : > 250 | Output Population : ef450+ Nuclei (T cell) |

| **Select Population (3)** | Input | Method | Output |
|---|---|---|---|
| | **Population :** Nuclei | **Method :** Filter by Property<br>Intensity ef450 Nuclei Mean : <= 250 | Output Population : ef450- (iPSC-aLO) |

| **Select Population (8)** | Input | Method | Output |
|---|---|---|---|
| | **Population :** ef450- (iPSC-aLO) | **Method :** Filter by Property<br>Intensity SPY555 Nuclei Mean : > 350 | Output Population : 555+ ef450- Nuclei (iPSC-aLO) |

| **Select Population (4)** | Input | Method | Output |
|---|---|---|---|
| | **Population :** 555+ ef450- Nuclei (iPSC-aLO) | **Method :** Filter by Property<br>Intensity AF647 Nuclei Mean : > 400 | Output Population : iPSC-aLO Casp pos |

| **Select Population (5)** | Input | Method | Output |
|---|---|---|---|
| | **Population :** 555+ ef450- Nuclei (iPSC-aLO) | **Method :** Filter by Property<br>Intensity AF647 Nuclei Mean : <= 400 | Output Population : iPSC-aLO Casp neg |

| **Select Population (6)** | Input | Method | Output |
|---|---|---|---|
| | **Population :** ef450+ Nuclei (T cell) | **Method :** Filter by Property<br>Intensity AF647 Nuclei Mean : > 400 | Output Population : T Cell Casp pos |

| **Select Population (7)** | Input | Method | Output |
|---|---|---|---|
| | **Population :** ef450+ Nuclei (T cell) | **Method :** Filter by Property<br>Intensity AF647 Nuclei Mean : <= 400 | Output Population : T cell Casp neg |

| **Calculate Properties** | Input | Method | Output |
|---|---|---|---|
| | **Population :** Organoid | **Method :** By Related Population<br>Related Population : Nuclei | Property Suffix : Number Nuclei Organoid |

| | | |
|---|---|---|
| | | Number of Nuclei |

| Calculate Properties (2) | Input | Method | Output |
|---|---|---|---|
| | **Population :** Organoid | **Method :** By Related Population<br>Related Population : iPSC-aLO Casp pos<br>Number of iPSC-aLO Casp pos | Property Suffix : per Organoid |

| Calculate Properties (3) | Input | Method | Output |
|---|---|---|---|
| | **Population :** Organoid | **Method :** By Related Population<br>Related Population : iPSC-aLO Casp neg<br>Number of iPSC-aLO Casp neg | Property Suffix : per Organoid |

| Calculate Properties (4) | Input | Method | Output |
|---|---|---|---|
| | **Population :** Organoid | **Method :** By Related Population<br>Related Population : T Cell Casp pos<br>Number of T Cell Casp pos | Property Suffix : per Organoid |

| Calculate Properties (5) | Input | Method | Output |
|---|---|---|---|
| | **Population :** Organoid | **Method :** By Related Population<br>Related Population : T cell Casp neg<br>Number of T cell Casp neg | Property Suffix : per Organoid |

| Calculate Properties (6) | Input | Method | Output |
|---|---|---|---|
| | **Population :** Organoid | **Method :** By Formula<br>Formula : A+B<br>Variable A : Number of T Cell Casp pos- per Organoid<br>Variable B : Number of T cell Casp neg- per Organoid | Output Property : Number T cells |

| Calculate Properties (9) | Input | Method | Output |
|---|---|---|---|
| | **Population :** Organoid | **Method :** By Formula<br>Formula : A+B<br>Variable A : Number of iPSC-aLO Casp pos- per Organoid | Output Property : Number iPSC-aLO cells |

| | | |
|---|---|---|
| | | Variable B : Number of iPSC-aLO Casp neg- per Organoid |

| Calculate Properties (7) | Input | Method | Output |
|---|---|---|---|
| | **Population :** Organoid | **Method :** By Formula<br>Formula : (A/(A+B))*100<br>Variable A : Number of iPSC-aLO Casp pos- per Organoid<br>Variable B : Number of iPSC-aLO Casp neg- per Organoid | Output Property : %Casp pos iPSC-aLO cells |

| Calculate Properties (8) | Input | Method | Output |
|---|---|---|---|
| | **Population :** Organoid | **Method :** By Formula<br>Formula : (A/(A+B))*100<br>Variable A : Number of T Cell Casp pos- per Organoid<br>Variable B : Number of T cell Casp neg- per Organoid | Output Property : %Casp pos T cells |

| Define Results | Results |
|---|---|
| | **Method :** List of Outputs<br>**Population : Image Region**<br>Number of Objects<br><br>**Population : Image Region 2**<br>Number of Objects<br><br>**Population : ef450+ Nuclei (T cell)**<br>Number of Objects<br><br>**Population : Nuclei**<br>Number of Objects<br><br>**Population : ef450- (iPSC-aLO)**<br>Number of Objects<br><br>**Population : 555+ ef450- Nuclei (iPSC-aLO)**<br>Number of Objects<br><br>**Population : iPSC-aLO Casp pos**<br>Number of Objects<br><br>**Population : iPSC-aLO Casp neg**<br>Number of Objects<br><br>**Population : T Cell Casp pos**<br>Number of Objects<br><br>**Population : T cell Casp neg**<br>Number of Objects<br><br>**Population : Organoid**<br>Number of Objects |

Number of Nuclei- Number Nuclei Organoid : Mean
Number of iPSC-aLO Casp pos- per Organoid : Mean
Number of iPSC-aLO Casp neg- per Organoid : Mean
Number of T Cell Casp pos- per Organoid : Mean
Number of T cell Casp neg- per Organoid : Mean
Number T cells : Mean
Number iPSC-aLO cells : Mean
%Casp pos iPSC-aLO cells : Mean
%Casp pos T cells : Mean

**Method :** Formula Output
Formula : a/b
Population Type : Objects
Variable a : Image Region - Number of Objects
Variable b : Image Region - Number of Objects
Output Name : Estimated No of Lung cells

**Object Results**
Population : Image Region : ALL
Population : Image Region 2 : ALL
Population : ef450+ Nuclei (T cell) : ALL
Population : Nuclei : ALL
Population : ef450- (iPSC-aLO) : ALL
Population : 555+ ef450- Nuclei (iPSC-aLO) : None
Population : iPSC-aLO Casp pos : ALL
Population : iPSC-aLO Casp neg : ALL
Population : T Cell Casp pos : ALL
Population : T cell Casp neg : ALL
Population : Organoid : ALL

Acapella version: 5.4.1.131587. Timestamp: 2025-05-07 11:08:23 +0200.
